# Live cell kinetic analysis of the LMO2/LDB1 leukemogenic protein complex reveals a hierarchy of turnover with implications for complex assembly

**DOI:** 10.1101/706259

**Authors:** Justin H. Layer, Michael Christy, Lindsay Placek, Derya Unutmaz, Yan Guo, Utpal P. Davé

## Abstract

Multisubunit protein complexes operate in many cellular functions. The LMO2/LDB1 macromolecular complex has been posited to be critical in hematopoietic stem and progenitor cell specification and in the development of acute leukemia. This complex is comprised of core subunits of LMO2 and LDB1 as well as bHLH and GATA transcription factors. We analyzed the steady state abundance and kinetic stability of LMO2 and its partners via Halo protein tagging in conjunction with variant proteins deficient in binding their respective direct protein partners. We discovered a hierarchy of protein stability, with half lives in descending order: LDB1>SSBP>LMO2>TAL1. Importantly, LDB1’s turnover was markedly prolonged and LDB1 conferred enhanced stability upon each and every subunit component thereby nucleating the formation of the multisubunit protein complex. Our studies provide significant insights into LMO2/LDB1 macromolecular protein complex assembly and stability, which has implications for understanding its role in blood cell formation and for therapeutically targeting this complex in human leukemias.

## Introduction

In hematopoiesis, lineage-specific transcription factors control specification of the hematopoietic stem cell (HSC) towards multiple diverse cell types. At the top of this developmental hierarchy are approximately 9 factors that directly affect the HSC itself: BMI1, RUNX1, GATA2, LMO2, TAL1, LDB1, MLL, GFI1, and ETV6 (Orkin and Zon, 2008). These master regulators are conserved among all vertebrates and have been experimentally characterized in mice, zebrafish, and humans (Jagannathan-Bogdan and Zon, 2013). The knockouts of any one of the genes encoding these factors causes the loss of all hematopoiesis, both embryonic and adult, by perturbing the creation, survival, or self-renewal of primitive and definitive HSCs. In examining this gene list, there are three emerging themes: First, the factors are part of a transcriptional network with autoregulation and inter-regulation (Wilson et al., 2010); second, the factors are frequently co-opted in human leukemias by various genetic mechanisms like chromosomal translocation (Greer, 2019); and, third, most remarkably for our study, all the factors function as part of multi-subunit protein complexes. Four of the factors listed above act in concert within a remarkable macromolecular complex, the LMO2/LDB1/TAL1/GATA2 (or the LDB1/LMO2) protein complex. There are diverse data supporting the idea that these proteins are bound together including co-immunoprecipitation (co-IP), co-purification followed by mass spectrometry, electrophoretic mobility shift assays, and co-occupancy at target genes by chromatin immunoprecipitation (Layer et al., 2016; Li et al., 2011; Meier et al., 2006; Wadman et al., 1997; Xu et al., 2003).

The assembly of the LDB1/LMO2 complex depends upon specific interactions between LMO2 and class II bHLH proteins, LMO2 and GATA factors, and LMO2 and LDB1. There are multiple bHLH and GATA paralogs capable of binding LMO2 so multiple versions of the LMO2-associated complex exist depending upon the expression of the subunits. LMO2 is an 18 kDa protein with two Zinc-binding LIM domains, LIM1 and LIM2. LIM1 folds to create an interface for binding class II bHLH proteins such as TAL1 and LYL1 (El Omari et al., 2011). LIM2 has an interface that binds GATA factors 1-3. A portion of LIM1 also serves as an interface for binding to the LIM interaction domain (LID) of LDB1. LDB1 has a self-association domain through which LDB1 may dimerize or multimerize (Liu and Dean, 2019). The class II bHLH proteins heterodimerize with class I bHLH proteins such as E2.2, E12, E47, and HEB (Murre, 2019). The bHLH proteins and GATA proteins can be part of the same complex allowing the LDB1/LMO2 complex to bind adjacent E boxes and GATA sites (Hewitt et al., 2016; Hewitt et al., 2015; Wadman et al., 1997; Xu et al., 2003). Such motifs bound by LMO2/LDB1 complexes have been described in erythroid progenitor cells at various gene targets including the beta globin gene promoters and the locus control region (LCR) (Hewitt et al., 2016; Li et al., 2011; Soler et al., 2010). The self-association domain of LDB1 mediates looping and proximity between the beta globin LCR and beta globin proximal promoters, a seminal example of enhancer-promoter communication (Deng et al., 2012; Krivega et al., 2014b; Liu and Dean, 2019; Song et al., 2007).

Several iterations of the LDB1/LMO2 complexes are drivers in leukemia. In fact, LMO2 and TAL1 were originally cloned from chromosomal translocations in T-cell acute lymphoblastic leukemia (T-ALL)(Nam and Rabbitts, 2006). LMO2 was also the target of insertional activation in gammaretroviral gene therapy-induced T-ALL (Davé et al., 2004; McCormack and Rabbitts, 2004). Mouse modeling and the characterization of the LMO2-associated complexes have been highly informative in dissecting the pathogenesis of LMO2-induced T-ALL, underscoring the role for specific bHLH and GATA factors as requisite co-operating drivers (Davé et al., 2009; McCormack et al., 2013; Ono et al., 1998; Smith et al., 2014). We recently confirmed by purification of FLAG-LDB1 and mass spectrometry that the LMO2/LDB1 complex in T-ALL closely resembles the complex hypothesized to function in normal HSCs (Layer et al., 2016).

Regardless of the variation in bHLH or GATA factors or the cofactors that these transcription factors may recruit, the core subunits of LMO2 and LDB1 are constant. We probed the LMO2/LDB1 interaction and discovered a discrete motif within the LDB1 LID that was essential for LMO2 binding. We consistently observed an increase in steady state abundance of LMO2 with co-expression of LDB1 and a decrease in abundance with the co-expression of LDB1ΔLID (Layer et al., 2016). Remarkably, this effect was observed in multiple leukemic cells including models for AML, which is consistent with recent studies showing the essentiality of LMO2 and LDB1 in these leukemias (Wang et al., 2017). To more closely analyze the effects on protein stability, we sought to understand the kinetics of turnover of LMO2 and its partner proteins. Towards this end, we devised a pulse chase technique through the use of multiplexed lentiviral expression of Halo-tagged proteins (Los et al., 2008). We discovered that there is a hierarchy of protein turnover for the subunits of the complex with LDB1 being the most stable protein. Furthermore, we discovered that every subunit, including both direct and indirect binding partners of LDB1, were stabilized by LDB1. These findings have remarkable implications for the assembly of this important macromolecular complex and underscore LDB1 as the major core subunit that could be targeted in leukemias.

## Results

### LMO2 turnover is mediated by ubiquitin-proteasomal system and is inhibited by LDB1

We first approached kinetic analysis of LMO2 turnover by quantitative western blotting after cycloheximide treatment. We observed half lives in the range of 8-10 hours for endogenous LMO2 in K562, MOLT4, and LOUCY leukemia cells; the half-life of exogenous LMO2 in Jurkat cells was measured at approximately 7 hours (data not shown). However, LDB1 decay was not observed by immunoblot within this same time frame. We were at the detection limits of our cycloheximide chase assay where cycloheximide toxicity is a confounding issue. Accordingly, we developed an alternative approach to analyze LMO2 and its associated proteins in live cells without metabolic perturbation and without toxins. We produced recombinant LMO2 tagged at its amino terminus with the Halo enzyme (Los et al., 2008). Our prior results showed that carboxyl terminal tags on LMO2 impeded its degradation so we focused on amino terminal tagging (Layer et al., 2016). We expressed Halo-LMO2 in Jurkat cells, which do not express endogenous LMO2, where the recombinant protein had enhanced steady state abundance with LDB1 co-expression (see lanes 6-7, Figure 1C), implying direct binding with LDB1. This was confirmed by co-immunoprecipitation (co-IP) of Halo-LMO2 with FLAG-LDB1 (data not shown). Confocal microscopy showed that Halo-LMO2 was localized predominantly in the nucleus (see Figures 1E-F). Thus, based on all of our conventional assays, Halo-LMO2 behaved just like untagged LMO2.

**Figure 1.**
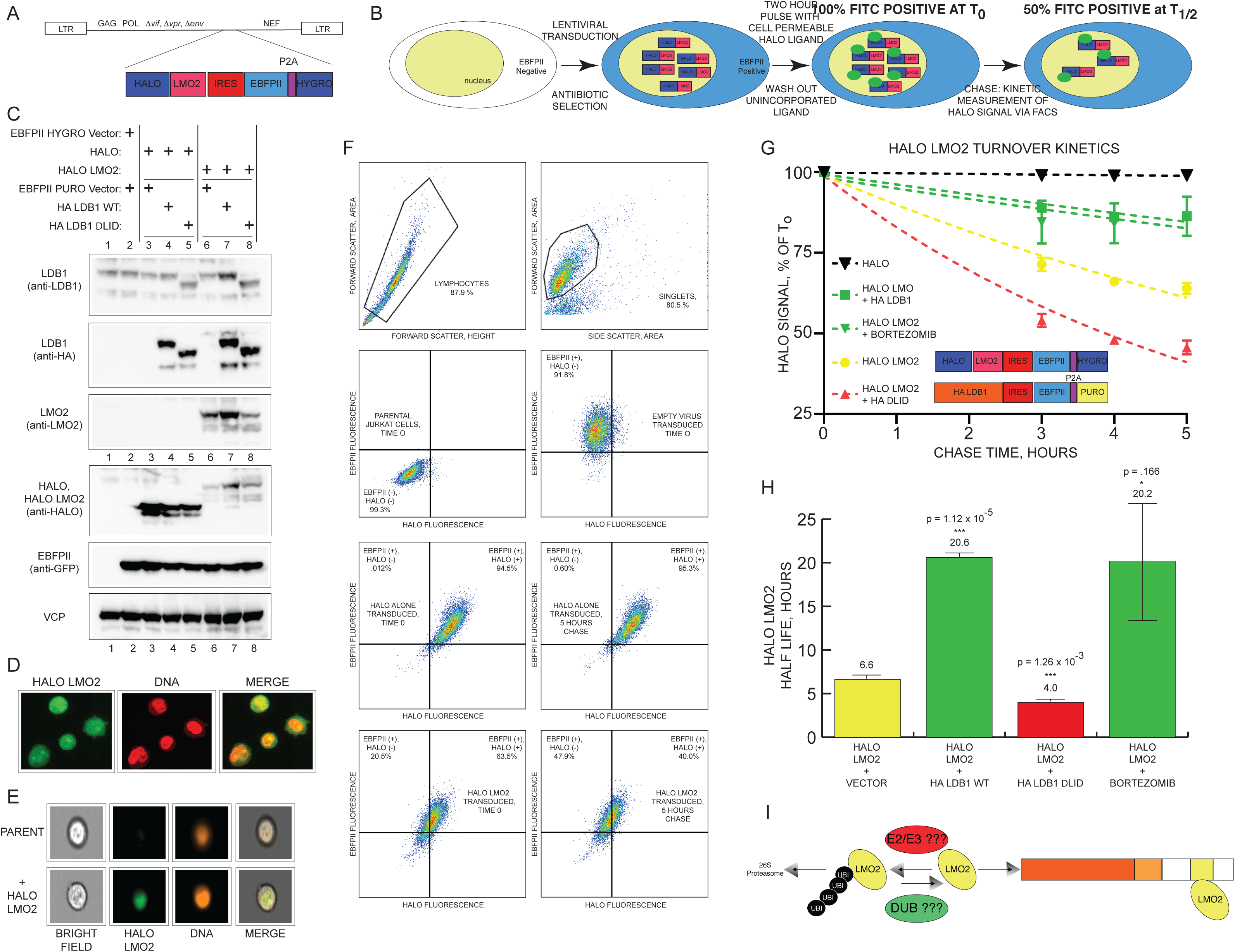
Pulse chase analysis of Halo-LMO2 in live cells demonstrates that LMO2 turnover is constrained by LDB1 and proteasomal inhibition. (A) Schematic showing the structure of the lentiviral expression vector; the recombinant expression cassette features a fluorescent protein and drug resistance proteins separated by a P2A protease site (see Materials and Methods). (B) Schematic showing the HaloLife assay. Cell transduction followed by pulse chase with cell-permeable Halo ligand. (C) SDS-PAGE immunoblot analysis of transduced cells. Expression (EBFPII) and loading controls (VCP) included. (D) Confocal microscopy images (E) Imagestream flow microscopy images (F) Flow histograms showing gating strategy for analysis of transduced cells. Bottom 3 histograms show EBFP fluorescence versus Halo fluorescence. Middle panels show untagged Halo protein; bottom panel shows Halo-LMO2 at t=0 (left) and t=5 h (right). (G) Plots of fluorescence decay during chase period. Curves were modeled to generate t_1/2_. (H) Bar graph showing the t1/2 of Halo-LMO2 with co-expression of LDB1, LDB1ΔLID, and bortezomib. (I) Model showing LMO2 stabilization by LDB1 when bound and degradation when unbound.

In order to force expression of multiple components of the LDB1/LMO2 complex in various cell lines individually and in combination, we developed multiplexed lentiviral expression vectors allowing fluorescence-based sorting and drug selection (Methods and Figure 1A and S1). Then, we implemented pulse chase analysis of Halo-tagged polypeptides by standard flow cytometry. We pulsed cells with the membrane-permeable fluorochrome, R110, and analyzed cellular fluorescence and R110 decay (i.e. chase) through the FITC channel throughout our experiments (Figure 1A-D). We called this technique for analyzing protein turnover, the **HaloLife** assay. As shown in Figure 1G, after a 90 min pulse of R110, we plotted the decay of fluorescence for untagged Halo protein and for Halo-LMO2 in the presence or absence of bortezomib, a specific 26S proteasomal inhibitor used in proteomic analysis of ubiquitinated moieties and also currently used to treat T-ALL (Kim et al., 2011; Raetz and Teachey, 2016). Bortezomib was tested with or without co-expression of HA-LDB1 or HA-LDB1ΔLID, which cannot bind LMO2. The curves fit a typical first order exponential decay, resulting in half-lives (t_1/2_) calculated and summarized in Figure 1H. Untagged Halo protein showed very slow protein turnover (Figure 1G), whereas Halo-LMO2 had a t_1/2_=6.6 hours, approximately the same t_1/2_ calculated from cycloheximide experiments. Co-expression of HA-LDB1 increased Halo-LMO2 t_1/2_ to 20.6 hours (P=1.12E-5). Similarly, bortezomib increased Halo-LMO2 t_1/2_ to 20.2 hours. In contrast, Halo-LMO2 was degraded faster with co-expression of HA-LDB1ΔLID (t_1/2_=4.0 hours, P= 1.26E-3). In summary, the presence of LDB1 markedly stabilized LMO2 as measured by the HaloLife assay. Halo-LMO2 turnover was reduced by bortezomib, implicating the ubiquitin-proteasomal pathway as the mechanism of degradation. Also, LDB1ΔLID, which is deficient in LMO2 binding but capable of homodimerization, increased the degradation of LMO2, a dominant negative effect which was previously observed in multiple leukemic cell lines (Layer et al., 2016).

### Specific LMO2 lysines are required for stabilization and are critical for binding to LDB1

The turnover of LMO2 is particularly intriguing since it is a known driver in T-cell leukemia and an essential factor in AML (Sun et al., 2013; Wang et al., 2017). Thus, the degradation of LMO2 could be exploited therapeutically to deplete the protein in diverse leukemias and lymphomas. Our prior experiments had discovered important features about the LMO2/LDB1 interaction: (1) binding is a prerequisite for LMO2 stabilization; (2) R^320^LITR within LDB1 are the key interacting residues and single residue substitutions within RLITR reduce LMO2 binding to LDB1; (3) I322 was accommodated by a hydrophobic pocket within LMO2 formed by L64 and L71 (Layer et al., 2016). Based on these data, we applied the HaloLife assay towards assessing the turnover of various mutant LMO2 proteins. Halo-LMO2(L64A, L71A) was significantly reduced in steady state abundance, and had faster turnover by measured t_1/2_=1.5 h compared to t_1/2_=6.2 h for Halo-LMO2 (Figure 2A-B). To identify the lysine residues within LMO2 that are potential sites for ubiquitination, we mutated the 10 lysines in the protein to arginine. Unexpectedly, lysine-less mutant LMO2 [denoted K(0)] had significantly faster turnover than LMO2 WT, t_1/2_=4.0 h versus 6.2 h(P=1.06E-3) (Figure 2B). We discovered that LMO2 K(0) was compromised in binding LDB1 as evidenced by reduced co-immunoprecipitation (Figure S2). We noted there were two lysines, K74 and K78, in proximity to the LMO2 hydrophobic binding pocket interfacing with LDB1 R^320^LITR. Halo-LMO2 (K74R, K78R), a mutant protein with only these two key lysines mutated and the remaining 8 lysines intact, showed significantly faster turnover, measured t_1/2_=3.9 h versus to t_1/2_ of Halo-LMO2 K(0) (P=1.76E-3). We also tested the reciprocal mutant, where we left K74 and K78 intact and mutated the remaining 8 lysines to arginine. As shown in Figure 2B, this mutant LMO2, Halo-LMO2 K(0)(K74, K78) had a measured t_1/2_=5.5 h, statistically insignificant (P=0.107) to the measured t_1/2_ of Halo-LMO2 WT. We then tested single substitutions at K74 and K78. Halo-LMO2 K(0)(K74) had a measured t_1/2_=4.8 h that was significantly (P7.28E-3) reduced compared to WT Halo-LMO2 whereas Halo-LMO2 K(0)(K78)’s t_1/2_ was not significantly different, t_1/2_=5.1 h (P=0.09) (Figure 2B). Intriguingly, K74 is conserved within all nuclear LIM-only proteins whereas K78 is unique to LMO2 (Figure 2D). Both K74 and K78 restored binding of the lysineless LMO2 to LDB1 (data not shown). Within lysineless proteins, the amino termini can serve as sites for ubiquitination. In order to show that the N-terminus of this version of LMO2 was critical for ubiquitin modification (Breitschopf et al., 1998; Trausch-Azar et al., 2004), we inserted a native LMO2 sequence translated from the longest transcript of the distal *LMO2* promoter, creating a super-stable protein, Halo-N+LMO2 K(0)(K74, K78) measured t_1/2_=25 h (P=4.47E-3). In summary, we identified K74 and K78 within LMO2 as essential for LDB1 binding and for normal levels of protein turnover.

**Figure 2.**
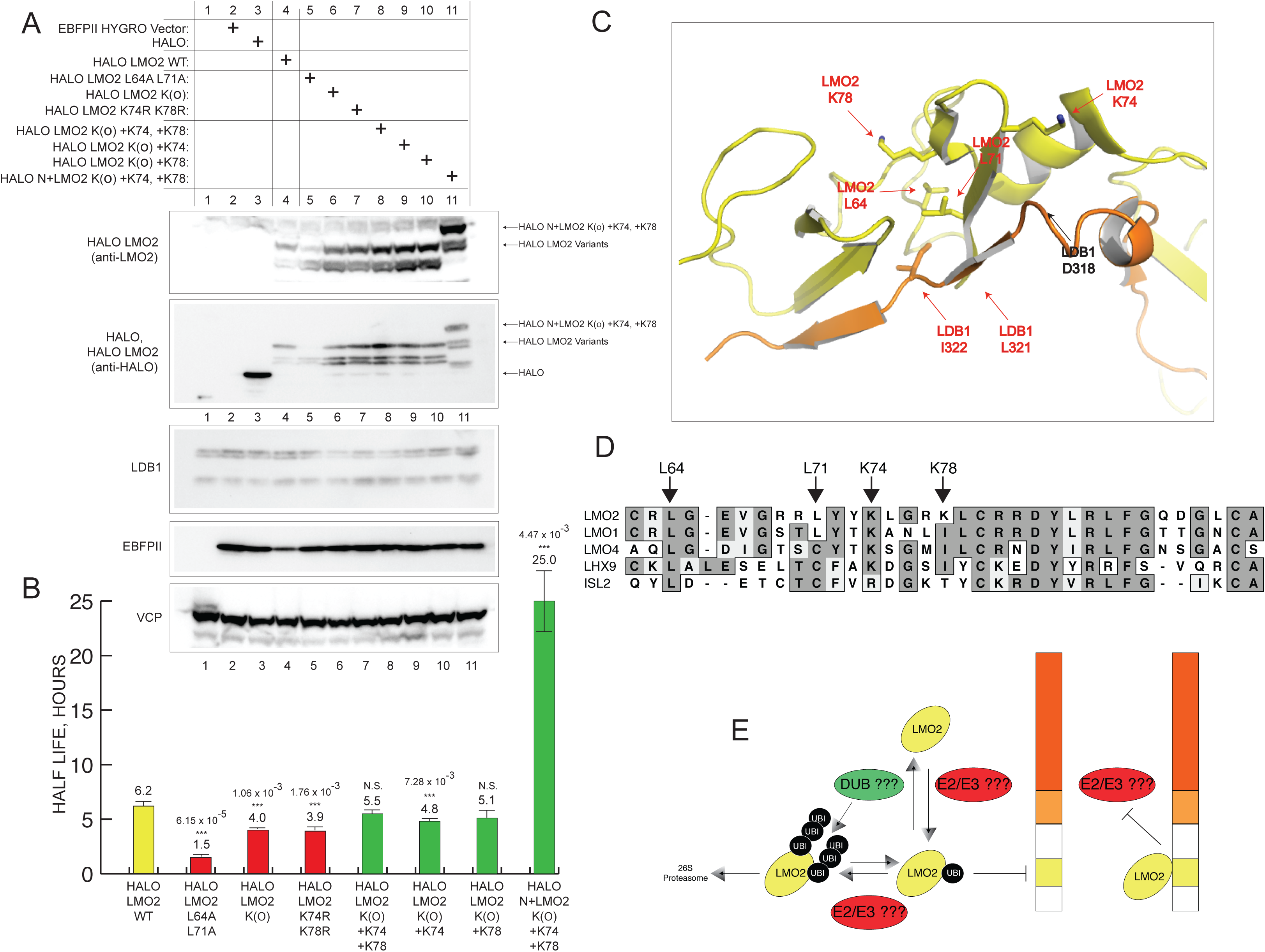
Critical Lysines K74 and K78 are required for LMO2/LDB1 binding and for LMO2 turnover. (A) Immunoblot analysis of various Halo-LMO2 proteins. Expression (EBFPII) and loading controls (VCP) included. (B) Half lives of Halo-LMO2 proteins and their variants. (C) PyMOL generated structure of the LMO2-LID fusion polypeptide. LMO2 backbone in orange and LID backbone in yellow. Key residues are discussed in text. (D) Alignment of LIM domain proteins. (E) Schematic showing a model for LMO2 stabilization by LDB1 and degradation in its free form.

Next, we examined the turnover of Halo-LMO2 in Jurkat, KOPT-K1, and K562 leukemia cells, which have various levels of LDB1 and LMO2. Jurkat cells are derived from T-ALL and express endogenous LMO1 but no LMO2; KOPT-K1 cells have a chromosomal translocation that results in overexpression of endogenous LMO2; and, K562 are aneuploid chronic myelogenous leukemia cells, resemble HSPCs, and express abundant endogenous LMO2 and LDB1 (Figure 3A) (Dong et al., 1995). Halo-LMO2 t_1/2_ was comparable in Jurkat and K562 cells, measured at 6.2 h versus 6.4 h, respectively. The super-stable Halo-N+LMO2 K(0)(K74, K78) was similarly prolonged, t_1/2_=25 and t_1/2_=20.9, respectively. In contrast, Halo-LMO2 t_1/2_ measured 1.3 h in KOPT-K1 cells. The fast turnover in KOPT-K1 cells suggested to us that forced expression of Halo-LMO2 was competing with high endogenous LMO2 (see lanes 5-8, Figure 3A) for the LDB1 LID. K562 cells had approximately equivalent abundance of LMO2 compared to KOPT-K1 cells, however, Halo-LMO2 turnover in K562 cells was not as fast perhaps due to the increased expression of endogenous LDB1 in comparison to KOPT-K1 cells (lanes 9-12, Figure 3A). Competition amongst LIM domain proteins is an important determinant of neuronal cell type specificity in the spinal cord. To test this competition model and its effect upon turnover, we measured Halo-LMO2 t_1/2_ and the effects of co-expression of competing nuclear LIM domain proteins: LMO2-HA, LMO1-HA, LMO4-HA, LHX9-HA, and ISL2-HA. These HA-tagged proteins expressed at various levels in Jurkat cells (lanes 4-8, Figure 4C) but their forced co-expression increased the turnover of Halo-LMO2 (Figure 3D). These results on t_1/2_ normalized to the level of expression achieved (Figure 3C), suggested an approximate order of affinity between LIM domain proteins for LDB1 LID. LMO2-HA was most competitive followed by LMO1, LMO4, LHX9, and ISL2. The LIM domain proteins that enhanced Halo-LMO2 turnover showed greater conservation of the key residues that we identified for LID binding, L64, L71, K74, and K78. All the LIM proteins tested had L64 conserved, however, only LMO1 and LMO2 have L71 (Figure 2D). LMO4 and LHX9 have a cysteine residue in place of K78 but have conserved K74 at the comparable position. Fitting this logic, ISL2, the protein that had no effect upon Halo-LMO2 turnover suggesting that ISL2 was the weakest competitor for LID binding, has an arginine residue in place of K74 and a threonine residue in place of K78.

**Figure 3.**
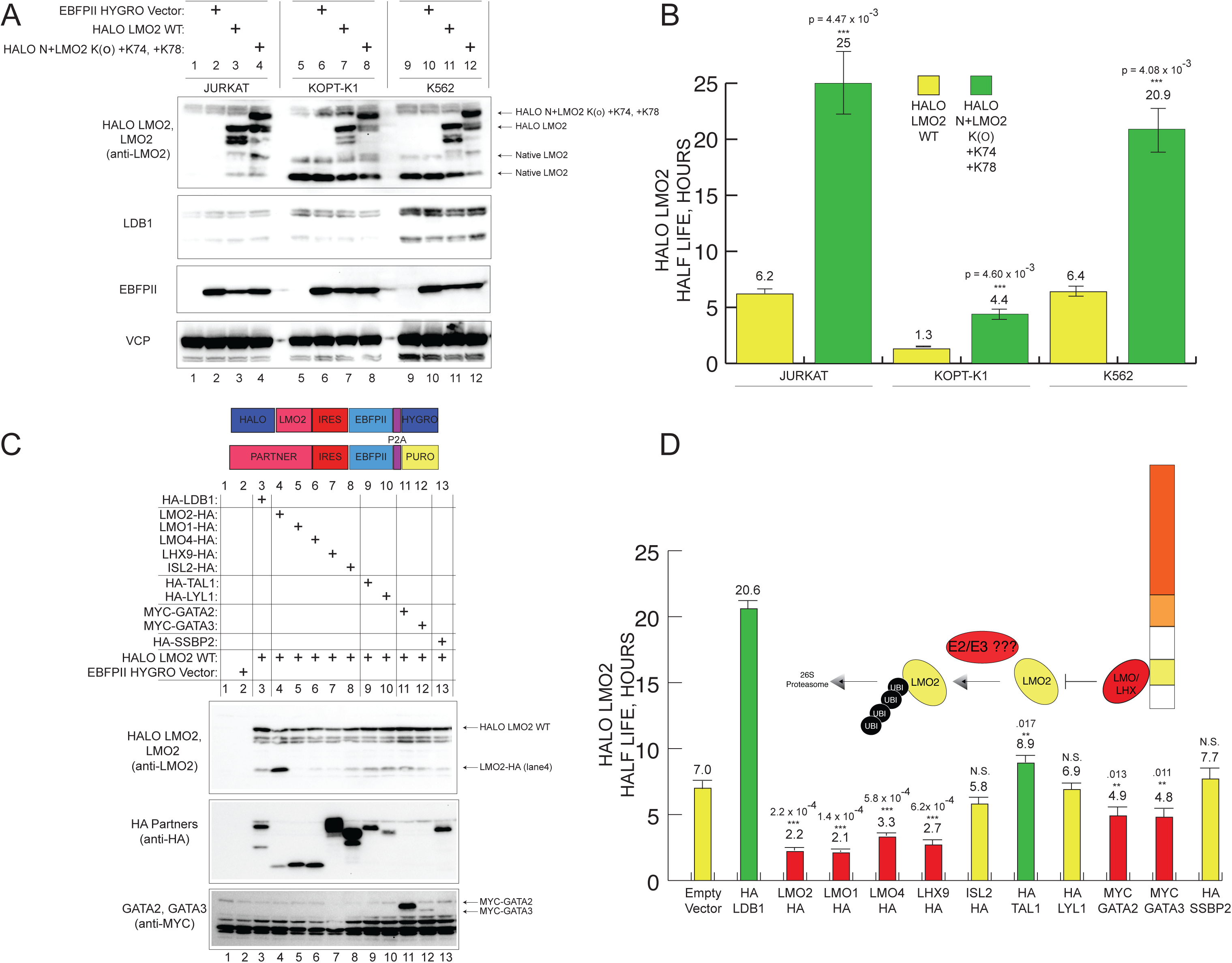
LIM domain proteins compete for LDB1 in leukemic cells and can accelerate LMO2 turnover. (A) Immunoblot showing Halo-LMO2 in various cell lines. Blots show endogenous LMO2 and LDB1 with expression and loading controls. (B) T1/2 for Halo-LMO2 and mutant Halo-LMO2 from HaloLife assay in Jurkat, KOPT-K1, and K562 cells. (C) Immunoblot of various HA-tagged LIM domain proteins transduced into Jurkat cells. (D) Bar graph showing half-lives of Halo-LMO2 with co-expression of various LIM domain proteins and other direct binding partners. P values for pairwise, two-tailed comparisons to empty vector are shown above the bars.

**Figure 4.**
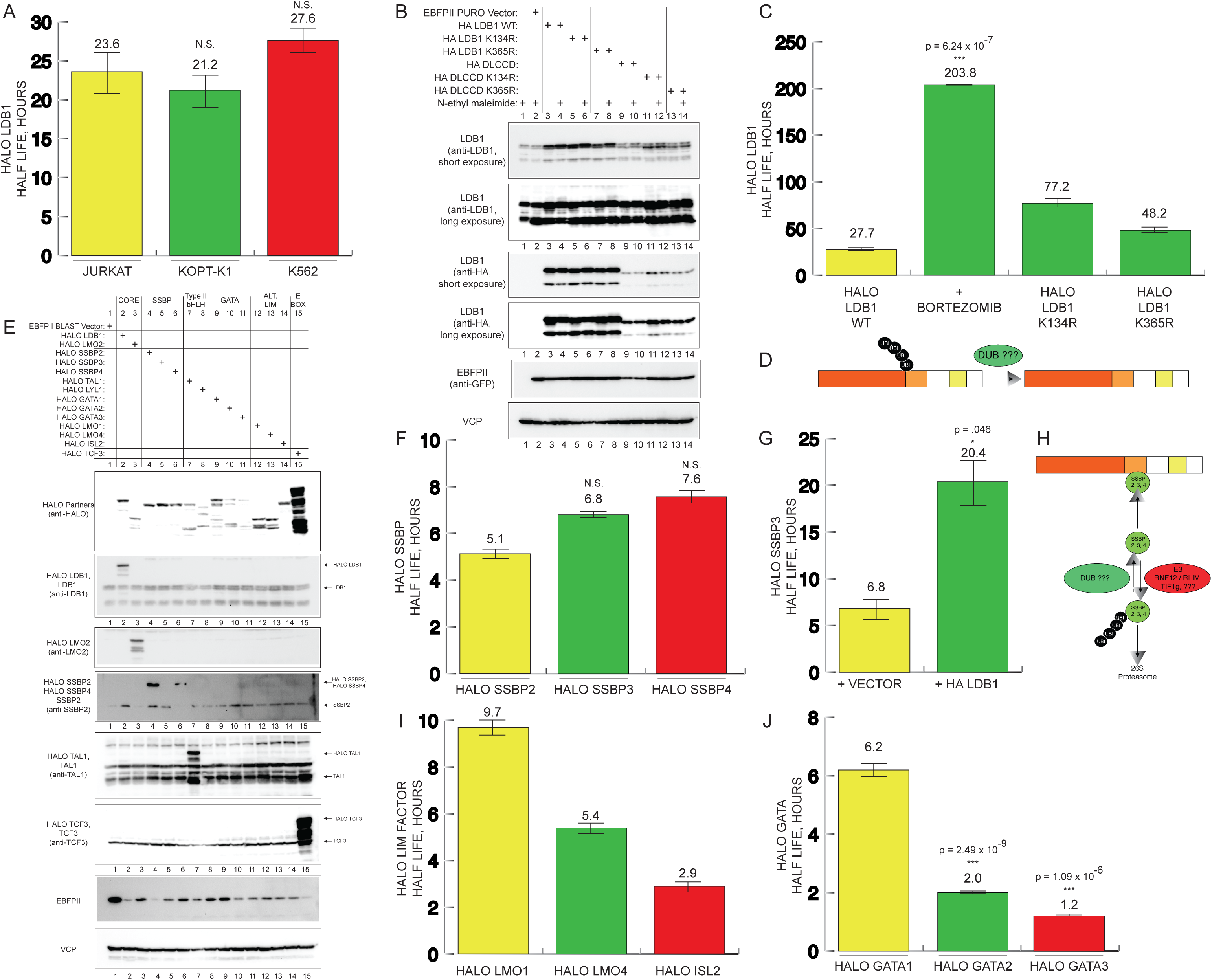
LDB1 is a long-lived protein in leukemia cells. (A) bar graph showing half lives of Halo-LDB1 in Jurkat, KOPT-K1, and K562 cells. (B) Immunoblot analysis of various Halo-tagged LDB1 proteins. All even lanes are extracts prepared in the presence of N-ethylmaleimide (NEM). (C) Bar graph showing half lives of Halo-LDB1, in the presence of bortezomib, and Halo-LDB1 K134R or Halo-LDB1 K365R. (D) Model showing ubiquitination on LDB1 K134. (E) Immunoblot analysis of Halo-LMO2, Halo-LDB1, Halo-TAL1, Halo-LYL1, Halo-SSBP2, and Halo-SSBP3. Expression and loading controls are shown. (F) Half lives of Halo-SSBP2, Halo-SSBP3, and Halo-SSBP4. (G) Half life of Halo-SSBP3 with vector and HA-LDB1 co-expression. (H) Schematic showing a model for SSBP degradation and stabilization by LDB1. (I) Half lives of Halo-GATA1, Halo-GATA2, and Halo-GATA3.

We also co-expressed other known LMO2 binding partners and measured their effects on LMO2 turnover. TAL1 increased Halo-LMO2 t_1/2_ to 8.9 h (P=0.017) but LYL1 did not change it from WT levels (6.9 v. 7.0 h, P=0.75). Co-expression of Myc-GATA2 and Myc-GATA3 both significantly decreased Halo-LMO2 to 4.9 (P=0.013) and 4.8 h (P=0.011), respectively. Myc-GATA3 expressed weakly but had a substantial effect on Halo-LMO2. Finally, Halo-LMO2 had a measured t_1/2_ of 7.7 h with HA-SSBP2 co-expression, a statistically insignificant change from WT turnover.

### LDB1 is a long-lived protein in leukemia cells

Based on the stabilization of LMO2, we suspected that LDB1 itself may be long lived and directly measured its turnover by Halo-tagging. Halo-LDB1 stability was consistent across diverse cell lines, measuring t_1/2_ of 23.6-27.6 h in Jurkat, KOPT-K1, and K562 cells (Figure 4A), respectively. Halo-LDB1 turnover was inhibited by bortezomib (Figure 4C). Prior studies had implicated K134 and K365 residues within LDB1 as affecting its degradation (Howard et al., 2010; Krivega et al., 2014a). Compared to LDB1 WT, which had t_1/2_ of 27.7 h, LDB1(K134R) and LDB1(K365R) half-lives were prolonged, t_1/2_=77.2 h and t_1/2_=48.2 h, respectively. Immunoblots of LDB1 showed two closely migrating bands, the slower band being enhanced in abundance with N-ethylmaleimide (NEM) (Figure 4B). This slower migrating band was not observed in blots for LDB1 (K134R) suggesting the addition of monoubiquitin at this residue.

In MEL and CHO cells, LDB1 stabilization was dependent upon Single Stranded DNA-Binding Protein 2 (SSBP2) (Xu et al., 2007). In contrast to these studies, LDB1 abundance did not increase with forced expression of SSBP2 or SSBP3 in any of the leukemic lines analyzed (data not shown). We directly tested the turnover of SSBP2 and SSBP3 by HaloLife analysis. Each paralog tested, SSBP2, SSBP3, and SSBP4, had faster turnover than LDB1, measured at t_1/2_=5.1 h and t_1/2_=6.8 h, and 7.6 h, respectively. SSBP2 and SSBP3 showed longer half-lives with LDB1 co-expression (Figure 4G). SSBP2 and SSBP3 stabilization was not seen with co-expression of LDB1ΔLCCD, the interaction domain between SSBP proteins and LDB1 (data not shown). However, the LDB1ΔLCCD mutant protein expressed at lower steady state abundance (see lanes 9-10, Figure 4B and S3), suggesting that there could be mutual folding and/or stabilization between SSBP proteins and LDB1. In summary, the HaloLife assay showed that every subunit of the LDB1/LMO2 complex had a shorter half-life than LDB1 and were subject to stabilization by LDB1.

### TAL1 and LYL1 are stabilized by the LMO2/LDB1 complex

TAL1 and LYL1 are necessary coöperating drivers in LMO2-induced leukemia (Ferrando et al., 2002; McCormack et al., 2013; Smith et al., 2014). These class II bHLH proteins are known binding partners of LMO2. The binding interface between TAL1 and LMO2 requires F238 within the second helix of the bHLH domain (Schlaeger et al., 2004), which is conserved as F201 within helix-2 of LYL1 (Figure 5A). We tested the turnover of Halo-TAL1 and Halo-LYL1 and specific mutants containing F238 and F201, respectively, by the HaloLife assay. Halo-TAL1 had a t_1/2_ of 4.2 h and Halo-LYL1 had a t_1/2_ of 1.8 h (Figure 5C, E). LMO2-HA co-expression did not significantly (t_1/2_=5.6 h with LMO2 v. t_1/2_=4.2 h without LMO2, P=0.215) stabilize TAL1 but stabilized LYL1 (t_1/2_=4.3 h v. 1.8 h, P=0.015). HA-LDB1 co-expression markedly stabilized Halo-TAL1 and Halo-LYL1 to t_1/2_ =19.9 h and t_1/2_ =20.5 h, respectively. This effect was only observed in the presence of LMO2. Similarly, Halo-TAL1 and Halo-LYL1 half-lives were similar to WT levels with co-expression of HA-LDB1ΔLID (Figure 5C, E). Thus, LDB1’s stabilization effect was not observed without LMO2 binding. To test the requirement for bHLH to LMO2 binding, we created mutant Halo proteins, Halo-TAL1(F238D), Halo-TAL1(F238G), Halo-LYL1(F201D), and LYL1(F201G), all of which were compromised in LMO2 binding in co-immunoprecipitation assays (data not shown). As expected, LMO2 did not stabilize these proteins. Each mutant bHLH protein had a measured t_1/2_ comparable to its WT counterpart. HA-LDB1 co-expression increased the t_1/2_ of Halo-TAL1(F238D) to 10.7 h (P=0.014). Similarly, Halo-LYL1(F201D) was stabilized by HA-LDB1 co-expression to t_1/2_ of 3.7 h (P=0.012). Thus, aspartic acid substitutions for F238 in TAL1 and F201 in LYL1 completely abrogated LMO2-induced stabilization but partially abrogated LDB1 induced stabilization. The F238D and F201D mutants may still retain some LMO2 binding especially since LDB1 stabilizes LMO2 and increases its steady state abundance. In contrast, glycine substitutions at the same residues completely abrogated both LMO2’s and LDB1’s effects. In summary, Halo-TAL1 and Halo-LYL1 half-lives in Jurkat cells, which are partially stabilized by LMO2 co-expression. Their half-lives are markedly prolonged by LDB1 co-expression but only if the proteins have intact LMO2 binding.

**Figure 5.**
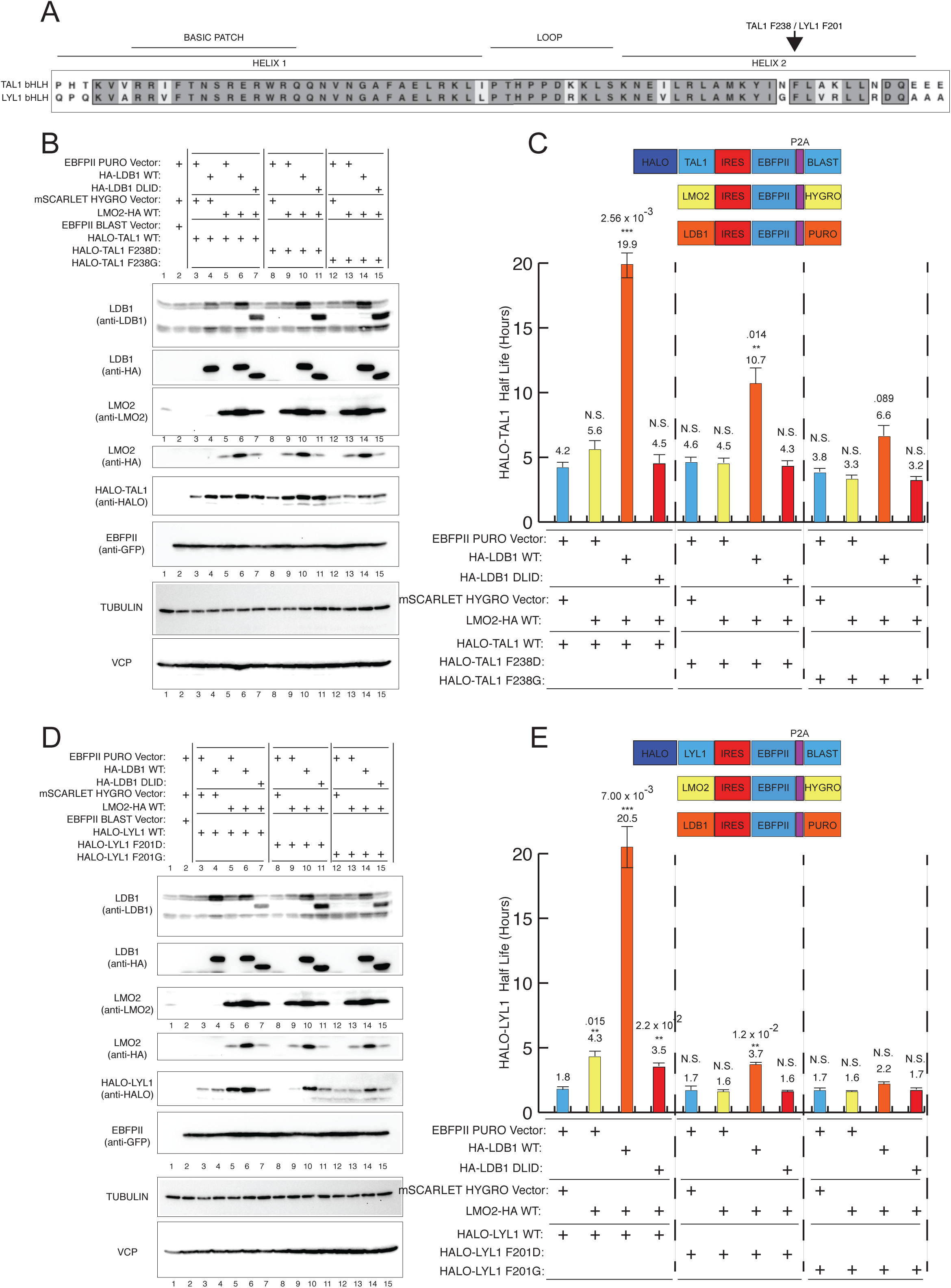
TAL1 and LYL1 are stabilized by LMO2/LDB1 binding. (A) Amino acid alignment of TAL1 and LYL1 bHLH domains. TAL1 F238 has been experimentally implicated in LMO2 binding corresponding to LYL1 F201. (B) Immunoblot of Halo-TAL1 or mutant TAL1 proteins expressed on their own or in the presence of LMO2-HA, HA-LDB1, or both. (C) Bar graph showing the half lives of Halo-TAL1 proteins in the absence or presence of LMO2-HA and HA-LDB1. Schematic above graph shows the expression cassettes with different antibiotic selection. (D) Immunoblot of Halo-LYL1 or mutant LYL1 proteins expressed on their own or in the presence of LMO2-HA, HA-LDB1, or both. (E) Bar graph showing the half lives of HALO-LYL1 proteins in the absence or presence of LMO2-HA and HA-LDB1. Schematic above graph shows the expression cassettes with different antibiotic selection.

### Complex assembly and function

Our results implied that intact binding interactions between all of the components created a stable macromolecular complex. We analyzed whether this assembly occurred in cells and whether complex assembly has a functional effect on transcription. Each component of our complex was expressed using a lentiviral vector with unique fluorescence and drug selection (Figures 1A and S1), We included empty vector controls (Figure 6A) as indicated. We transduced components pairwise with or without FLAG-LDB1 (F-LDB1) to test abundance (Figure 6A) and binding (Figure 6B) by co-immunoprecipitation with anti-FLAG monoclonal antibody. The measured half-lives uniformly explained increased steady state abundances of Halo-tagged proteins detected by Western blot analysis. The experiments in Figure 6A extend this correlation to untagged or minimally tagged (i.e. single HA) proteins as well. SSBP2 was poorly expressed in Jurkat cells so SSBP3 was transduced instead; our prior experiments had shown comparable peptide counts for SSBP3 and SSBP2 by tandem mass spectrometry of purified LDB1 complexes (Layer et al., 2016). HA-SSBP3 was stabilized by LDB1 but not by co-expression of LMO2 (see lanes 6-9, Figure 6A). Consistent with the HaloLife results, TAL1 and LYL1 were maximally stabilized by the co-expression of both LMO2 and LDB1 (see lanes 10, 11 to 12, 13 for TAL1 and lanes 18, 19 to 20, 21 for LYL1).

**Figure 6.**
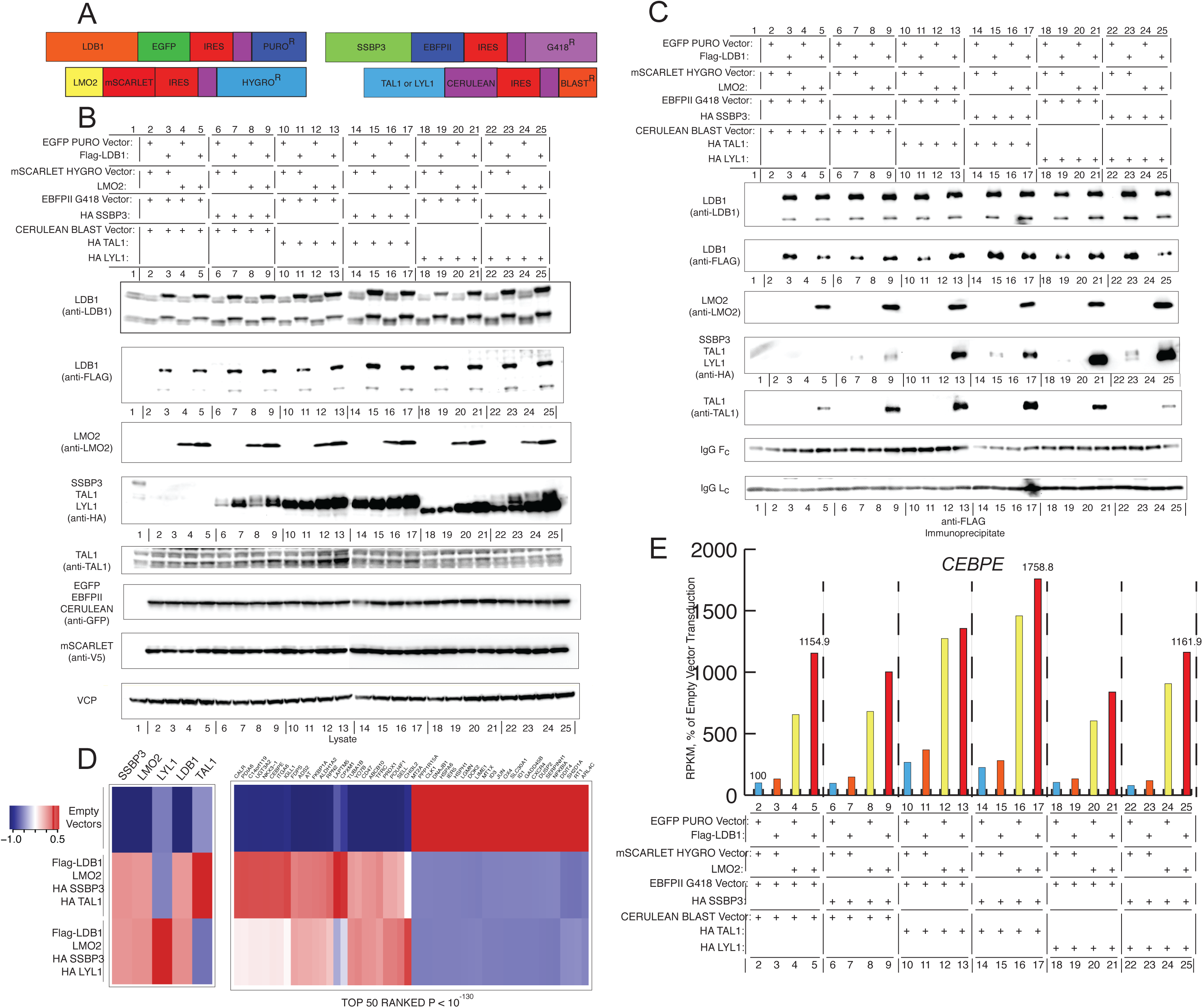
Reconstitution of the LMO2/LDB1 complex and its transcriptional output. (A) Schematic showing the lentiviral expression cassettes with fluorescent protein expression and antibiotic selection. (B) Immunoblot analysis of whole cell lysates prepared from Jurkat cells transduced with the respective proteins. Expression control is shown by anti-GFP or anti-V5 in the case of mScarlet. Two independent loading controls, anti-tubulin and anti-VCP, are shown. (C) Immunoblots of immunoprecipitations of Flag-LDB1 with anti-Flag. (D) Heat map showing the top 50 genes and their expression in 3 different transduction groups, empty vectors, LMO2/LDB1/SSBP3/TAL1, and LMO2/LDB1/SSBP3/LYL1.

Complex assembly was analyzed by anti-FLAG immunoprecipitation via F-LDB1. Jurkat cells have abundant endogenous TAL1, which was immunoprecipitated by F-LDB1 only in the presence of LMO2 (lanes 2-5, Figure 6B). Endogenous TAL1 co-IP was augmented by co-expression of SSBP3 (lanes 6-9, Figure 6B). Forced expression of LYL1 did not effectively outcompete endogenous TAL1 for LMO2/LDB1 binding whereas SSBP3 and LYL1 co-expression reduced steady state TAL1 and TAL1 co-IP (see lanes 21 and 25, Figure 6B). Next, we analyzed the effects of complex formation upon gene expression. We performed a pairwise comparison of RNA-seq on Jurkat cells transduced with all complex components (i.e. LMO2, LDB1, SSBP3, and TAL1 or LYL1; lanes 17 and 25 in Figure 6) versus cells transduced with empty virus (lane 2, Figure 6), generating a ranked list of differentially expressed genes. Most of the genes on this list were maximally activated or repressed by co-expression of the full complex and not by expression of partial complex components, as shown for activation of ALDH1A2, CEBPE (Figure 6E), and NKX31, and other bona fide targets (Figure 6D).

### HaloLife assay can be used to screen for modifiers of degradation

Next, we asked whether the stable leukemia lines expressing various Halo-tagged proteins can be used in a screen to identify modifiers of stability. Deubiquitinases (DUBs) of the LMO2-associated proteins would stabilize LMO2 complex formation and could be important therapeutic targets in leukemias dependent upon LMO2. Also, the number of genes encoding DUBs was suitable for a targeted screen, ∼80 genes versus ∼400 genes encoding E3 enzymes (Komander and Rape, 2012). We assembled a lentiviral shRNA library against 70 DUB genes, of which 44 (63%) were expressed in Jurkat cells. We transduced pooled shRNAs directed against each DUB into individual Jurkat lines stably expressing Halo-LMO2, Halo-LDB1, Halo-SSBP2, Halo-SSBP3, Halo-TAL1, or Halo-LYL1. After transduction, we analyzed the cells for their growth and for effects on the Halo-tagged proteins. We devised three criteria to identify an important hit: (1) if the percentage of R110 fluorescence was reduced at t_0_ in cells transduced with a DUB-specific shRNA compared to scrambled shRNA; (2) a reduction in absolute Halo signal (i.e. MFI) at t_0_; or, (3) a reduction in Halo signal after a 5 h chase (Figure 7A). Figures 7B and S show the outcomes of this screen. We identified a set of shRNAs against a DUB, ALG13, that met all 3 criteria for every subunit of the complex: Halo-LMO2, Halo-LDB1, Halo-SSBP2, and Halo-SSBP3 and 2 criteria for Halo-TAL1 and Halo-LYL1 (Figure 7B). Other DUBs that potentially affected some of the subunits met 2 out of 3 criteria including OTUD7B, USP3, and USP4 (Figure S4). ALG13 is a DUB with an unusual structure. ALG13 has an amino-terminal glycosyltransferase domain (Gao et al., 2005) followed by the DUB domain found in the Ovarian Tumor (OTU) class of DUBs and a tudor domain followed by a proline rich domain (Mevissen et al., 2013). The OTU family of DUBs had several hits meeting our criteria for various subunits (Figure 7B). The pool of shRNAs against ALG13, was validated in a secondary screen and a time course for Halo-LMO2 degradation (Figure 7C). As shown in Figure 7C, the ALG13 shRNA knockdown accelerated the degradation of Halo-LMO2 compared to transduction of scrambled shRNA control or shRNAs directed against an OTU DUB that is not expressed in Jurkat cells (OTUB1). The ALG13 shRNA pool was comprised of 5 shRNAs, which we tested individually in the same assay. Four out of the 5 shRNAs caused increased turnover of Halo-LMO2 (data not shown). To further validate the role of ALG13 in LMO2 degradation, we performed the HaloLife assay by forcing the expression of full length ALG13 (1137 aa) or catalytically inactive mutant ALG13 and measuring the resultant t_1/2_. We deleted the DUB domain creating ALG13ΔDUB (deleted catalytic DUB domain) but could not rule out drastic effects upon folding of the protein so we engineered a point mutant, ALG13 C242R. Interestingly, alanine substitution at the catalytic cysteine residue can enhance the affinity for ubiquitin in OTU DUBs so an arginine substitution is the better residue to evaluate a catalytically inactive DUB (Morrow et al., 2018). We measured t_1/2_ of Halo-LMO2 of 6.4 h in empty vector control but with forced expression of full length ALG13, we measured t_1/2_=7.6 h (P=0.009 for comparison to empty vector control). In contrast, we measured t_1/2_= 6.7 h (P=NS) and 6.3 h (P=NS) with ALG13ΔDUB and ALG13 C242R mutant proteins, respectively.

**Figure 7.**
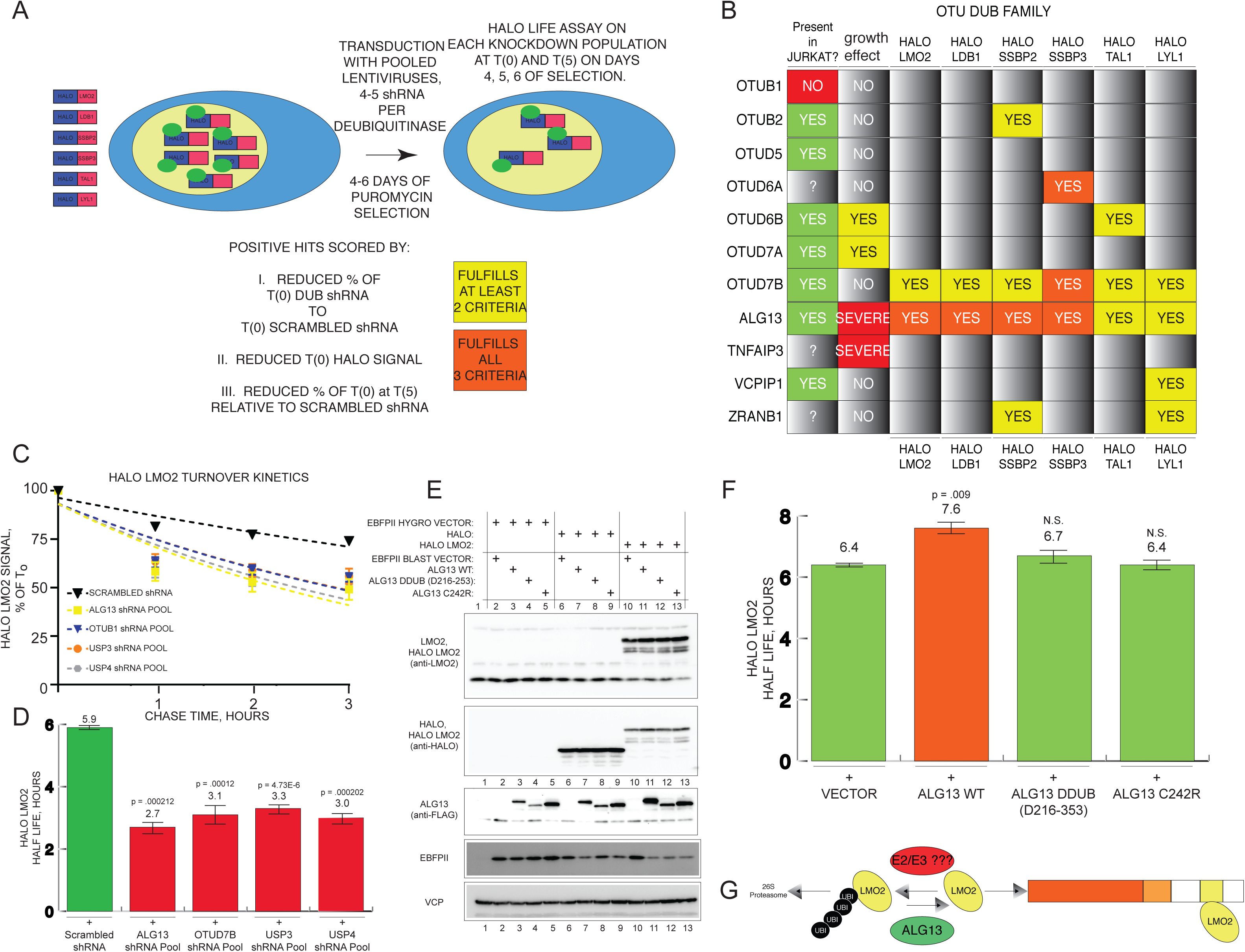
HaloLife screen of DUB genes. (A) Schematic shows the experimental assay for shRNA screening for DUBs. Yellow denotes DUB shRNA knockdowns that fulfilled 2 of the 3 stated criteria whereas red denotes those knockdowns that fulfilled all 3 criteria. (B) Table shows hits within the OTU DUB family of genes. (C) Decay curve of Halo-LMO2 after shRNA knockdown of respective DUB RNAs. (D) Immunoblot of FLAG-ALG13 proteins, WT, ΔDUB, or C242R in K562 cells. (E) Half lives of Halo-LMO2 with co-expression of vector, or ALG13 WT, ALG13ΔDUB, or ALG13(C242R).

## Discussion

In this study, we describe a novel technique to analyze the turnover of the components of the leukemogenic LMO2/LDB1 protein complex, employing Halo-tagging and fluorescence-based pulse chase analysis. The assay, which we termed HaloLife, is informative in that the turnover of tagged proteins is observed in live cells. Thus, proteins are observed in their natural milieu without pharmacologic, nutritional, or mechanical disruption. This method has the added advantage of allowing the testing of the effects of various culture conditions and small molecule therapeutics upon protein turnover. The Halo tag is advantageous because it is relatively small and monomeric, approximately the mass of GFP, which has been used in similar studies. Of course, as is the case in all epitope tagging, one must verify that the tag itself does not disrupt the behavior of the protein. In the case of the proteins presented here, each one was localized to the nucleus (Figure S5) and retained its affinity for its physiologic partners. Also, mutations that disrupted binding had the same effect upon Halo-tagged versions as the untagged proteins themselves. The pulse chase analysis showed that the Halo protein itself was very long lived (t_1/2_>100 h). Each Halo-tagged protein had rapid turnover compared to Halo itself, such that the fusion proteins acted as “degrons” for the Halo protein. In light of the caveats noted, the t_1/2_ measured in the HaloLife assay can be viewed as an approximation of the true half-life of the native protein. However, all the measured half-lives in this study closely matched those estimated from cycloheximide chase and quantitative immunoblotting (Lurie et al., 2008) and provided an explanation for detected changes in steady state abundance. In summary, the HaloLife has the compelling advantages of being performed in live cells, in their native cellular milieu, and at steady state without cellular disruption.

HaloLife analysis of LMO2 and its binding partners revealed a hierarchy of protein turnover with LDB1 being the most stable protein. Observed half-lives in Jurkat cells in increasing order were: Halo-LYL1 (∼1.8 h), Halo-TAL1 (∼4.1 h), Halo-LMO2 (∼6.4 h), Halo-SSBP2 (∼5.1 h), Halo-SSBP3 (∼6.8 h), and Halo-LDB1 (∼20-24 h). Most remarkably, co-expression of LDB1 shifted the turnover of these Halo tagged subunits so that each protein partner assumed a half-life of ∼20 h in the presence of excess LDB1, approximating the measured half-life of LDB1 itself. There was no reciprocal effect since none of the partner proteins prolonged the half-life of LDB1. All proteins tested were markedly stabilized by bortezomib, suggesting degradation by the ubiquitin proteasomal system. Each protein partner had to bind to LDB1 either directly or indirectly, in the case of TAL1 and LYL1, to be stabilized. Taken together, these findings suggest that the free subunits, those unbound to LDB1, are degraded more rapidly than those bound to LDB1. Furthermore, the prolonged half-life of LDB1 suggests that it is the core subunit in the assembly of the bHLH/LMO2/SSBP/LDB1 macromolecular complex, which we term the LDB1/LMO2 holocomplex. As LDB1 binds to its direct partners, SSBP proteins or LMO2, LDB1 impedes the turnover of other components of the complex so that stepwise assembly and slow turnover increase the steady state abundance of the holocomplex. Accordingly, each subunit assumes a half-life similar to that of LDB1, suggesting that the whole complex may be degraded *en masse*. Two distinct lysines within LDB1, K134 and K365, have been implicated in LDB1 turnover. Both K134R and K365R mutations markedly prolonged LDB1 turnover by the HaloLife assay compared to wild type LDB1, thereby confirming the role of these lysine residues in LDB1 stability. Neither lysine is within a domain mediating subunit binding (i.e. LDB1’s LCCD, residues 200-249, is responsible for SSBP binding and the LID is comprised of residues 300-330), Thus, these residues are unlikely to be occluded from ubiquitination by SSBP or LMO proteins. On the other hand, K134 is within the dimerization domain, so K134 could be masked by homodimerization. This raises the possibility of LDB1 homodimers being more stable than monomers. We discovered a slower migrating LDB1 in the presence of N-ethylmaleimide that is consistent with a monoubiquitin conjugation to K134. If we assume this residue is only accessible in unbound LDB1, then we predict that this monoubiquitinated LDB1 is monomeric. Although the stoichiometry of the LDB1 holocomplex has not been definitively solved, our prior mass spectrometry data do suggest stable LDB1 dimers in nuclear lysates. Interestingly, this theme of accessible lysines may be extended to the turnover of LMO2 and SSBP proteins as well. Our experiments with LMO2 implicated K74 and K78 in LDB1 binding. These residues may be sites of ubiquitination and may be exposed in free LMO2 subunits but sterically hindered in LMO2 bound to LDB1. Alternatively, K74 and K78 may be subject to other post-translational modifications such as methylation or acetylation. K78 is particularly intriguing since it is unique to LMO2 and is adjacent to a hydrophobic pocket (L64 and L71) such that neutralization of the side chain amine would favor LDB1 binding by accommodating I322. This contact interface is supported by a crystal structure of an LMO2-LID fusion protein (El Omari et al., 2011). We co-purified SSBP3 with FLAG-LDB1 and detected a diGly motif on K35 in the mass spectrometry data (data not shown), which could be a remnant of trypsinized ubiquitin, although NEDD8 and ISG13 are other possible conjugates (Emanuele et al., 2011). Nevertheless, K35, K7, and other conserved lysines are within the LUFS domain of SSBP proteins and are expected to be masked by LDB1 binding whereas free SSBP subunits should have more accessible lysine residues for modification. In summary, free subunits of the LMO2/LDB1 complex are rapidly degraded in comparison to the slow degradation kinetics of the holocomplex. Complex assembly may proceed through binding and stabilization by masking key lysine residues in the free subunits. Recombinant full-length proteins and a structure of the holocomplex may be able to test this model. On a more general note, our studies suggest that multisubunit protein complexes may have key core subunits with enhanced stability that can be conferred upon binding subunits. To name a few examples, core subunits analogous to LDB1 exist for the T-cell receptor, BAF complex, Mediator complex, and TFIID protein complexes (Bonifacino et al., 1990; Cai et al., 2010; Imasaki et al., 2011; Mashtalir et al., 2018; Wright et al., 2006). It would be interesting to see whether lysine residues targeted for ubiquitination are masked in other macromolecular assemblies as well.

Prolonged turnover of nuclear factors and transcription factors has been suggested to be due to their association with chromatin. The subunits of the LDB1/LMO2 complex were localized to the nucleus, at least 2-fold over cytoplasm but we could not analyze whether they were chromatin-bound. The slow turnover of the LMO2/LDB1 holocomplex obviates the need to form new chromosomal loops that co-localize enhancers to core promoters during every cycle of RNA Pol II recruitment, which would be energetically unfavorable. Notably, co-expression of all complex components resulted in maximal target gene activation or repression implying that assembly of the holocomplex is what is needed to effect gene regulation.

It is important to note that the HaloLife assays were all performed in leukemic cells. The leukemia lines were of diverse lineages. Even so, one cannot rule out a general defect in the turnover of LMO2 and LDB1 in all of these lines. The work shown here required the development of novel lentiviral vectors to allow co-expression of all complex partners in the same cell. Similar analysis in normal hematopoietic cells would be challenging but is being explored since the turnover and stoichiometry of this complex in primary hematopoietic cells is of great interest and a part of our ongoing research. Lentiviral transduction of hematopoietic stem cells is inefficient and co-expression by multiple transductions would be very challenging. Of course, studying the turnover of LMO2 and LDB1 in leukemic lines is suitable for studying leukemia pathogenesis. Importantly, careful analysis of this protein complex turnover has major implications for regulating these major drivers of leukemia. Recent data from mouse genetics strongly supports a role for Ldb1 in Lmo2-induced leukemia. The *CD2-Lmo2* transgenic mouse model develops T-ALL with long latency but with complete penetrance (Smith et al., 2014). Conditional deletion of *Ldb1* in this model abrogated T-ALL onset (UPD personal observation). Thus, Ldb1 is a required Lmo2 partner in this murine model of T-ALL. This compelling result from mouse genetics coupled with the primacy of LDB1 in a protein turnover hierarchy underscore the potential for targeting the LMO2/LDB1 interface in leukemias. If LMO2 is dissociated from LDB1 then free LMO2 and TAL1 are expected to undergo rapid degradation. Supporting this idea, the co-expression of LIM domain proteins that competed for the LID (LMO1, LMO2, LMO4, and LHX9) accelerated Halo-LMO2 turnover. ISL2, which has the least similarity to LMO2 residues responsible for LID binding, did not accelerate turnover, underscoring the determinants of LID binding as a mechanism for LIM protein competition. We predict a small molecule that could bind to the LID interface would also accelerate LMO2 turnover. Of course, such an inhibitor of LMO2 binding to LDB1 would affect normal hematopoietic stem cells as well. However, there could be a therapeutic index with higher LMO2/LDB1 holocomplex-expressing cells predicted to be more sensitive to such inhibition.

Previous work implicated RNF12 as a potential E3 enzyme responsible for LDB1 and LMO2 degradation (Güngör et al., 2007; Ostendorff et al., 2002; Xu et al., 2007). However, in our experiments, steady state abundance of LDB1 and other subunit proteins were unchanged with forced expression of RNF12 in Jurkat cells (data not shown). Thus, additional investigation is needed to characterize the degradation machinery of the LMO2 holocomplex especially in its normal or leukemic cellular contexts, which could reveal E3 enzymes or DUBs that could be therapeutically targeted. DUB enzymes are particularly amenable to small molecule inhibition since proteolytic mechanisms have been extensively studied. An shRNA knockdown screen using the HaloLife assay showed a very compelling candidate DUB, ALG13. There were other candidates identified in our screen such as OTUD7B, but ALG13 fulfilled our screening criteria and affected all subunits with no effect upon Halo protein itself. Recently, with the development of Proteolysis Targeting Chimeras (i.e. PROTACs), there is great interest in small molecules that can induce targeted degradation by recruitment of E3s to proteins of interest (Deshaies, 2015). Actually, one of these PROTACs is being analyzed in phase II clinical trials with similar molecules on the horizon (Lai and Crews, 2017). In contrast, bortezomib is being tested in a randomized clinical trial in T-ALL as an addition to state of the art multiagent chemotherapy. The results from our study show that bortezomib stabilizes LMO2 oncoprotein, which can potentially antagonize the effect of chemotherapies. However, the overall effect of bortezomib upon T-ALL and patient survival are difficult to predict since bortezomib affects pathways other than LMO2 causing proteotoxic stress in leukemic cells (Vilimas et al., 2007). Our ongoing work on LMO2/LDB1 complex turnover should be highly revealing for both normal hematopoietic stem cell biology and for the development of novel leukemia therapies.

## Acknowledgements

We thank Drs. Yuichiro Takagi, Sabine Wenzel, Mark Goebl, Merv Yoder, Reuben Kapur, and Jörg Bungert for helpful discussions. This work was supported in part by Merit Review Award # I01BX001799 from the United States Department of Veterans Affairs, Biomedical Laboratory Research and Development Service, and R01CA207530 from the National Cancer Institute, awarded to UPD. Additional support was provided to UPD by the Strategic Research Initiative of the Indiana University School of Medicine. JHL was a recipient of the Biomedical Research Grant from Indiana University. We acknowledge the IUSCC Flow Cytometry Core and the Genomics Core. JHL and UPD both thank Drs. Steve Brandt, Mark Koury and Ray Mernaugh, for all of their guidance and help through the years. JHL acknowledges and thanks Dr. P.A. Weil, for consistently imparting the value and power of thoughtful experimental controls, amongst every one of his many trainees.

## MATERIALS AND METHODS

### CONTACT FOR REAGENT AND RESOURCE SHARING

Further information and requests for resources and reagents should be directed to and will be fulfilled by the Lead Contact, Dr. Utpal Davé.

| REAGENT or RESOURCE | SOURCE | IDENTIFIER |
| --- | --- | --- |
| Antibodies |  |  |
| mouse monoclonal antiLDB1 IgG | Santa Cruz | Cat# sc-376030x |
| goat antimouse IgG Fc-horseradish peroxidase (HRP)conjugate | Thermo/Pierce | Cat#31439 |
| mouse antivalosin-containing protein antibody | Abcam | Cat#ab11433 |
| mouse Anti LMO2 | Levy, Layer |  |
| anti FLAG-HRP conjugate | Sigma | Cat#A8592 |
| anti HA-HRP conjugate | Roche | Cat#12013819001 |
| anti V5-HRP conjugate | Invitrogen | Cat#46-0708 |
| rabbit polyclonal anti TAL1 IgG | Bethyl | Cat#A305-300A |
| goat anti rabbit IgG conjugate | Jackson ImmunoResearch | Cat#211-032-171 |
| mouse monoclonal anti SSBP2 IgG | Santa Cruz | Cat#166687 |
| mouse monoclonal anti HALO IgG | Promega | Cat#G921A |
| mouse monoclonal anti GFP IgG | Roche | Cat#11814460001 |
| Rabbit polyclonal anti tubulin IgG | Santa Cruz | Cat#SC9104 |
| mouse anti valosin-contaning protein (VCP) | Abcam | Cat#ab11433 |
| Chemicals, Peptides, and Recombinant Proteins |  |  |
| Bortezomib | MP Biomedicals, LLC | Cat#180869 |
| HaloTag Ligand R110 | Promega | Cat#G3221 |
| SYTO 17 red fluorescent nucleic acid stain | Molecular Probes Inc. | Cat#S7579 |
| pBluescript SK | Stratagene |  |
| Iscove's modified Dulbecco's medium<br>(IMDM) | Gibco | Cat#12200-036 |
| RPMI 1640 | Gibco | Cat#31800-022 |
| Penicillin Streptomycin Solution 10X | Corning | Cat#30-022-CI |
| Geneticin | Gibco | Cat#10131-027 |
| 0.05% Trypsin, 0.53 mM EDTA 1X [-]sodium<br>bicarbonate | Corning | Cat#20116004 |
| Puromycin dihydrochloride | Fisher Bioreagents | Cat#BP2956-100 |
| Pierce Protease Inhibitor Tablets | ThermoScientific | Cat#A32965 |
| Hygromycin B in PBS 50mg/ml | Invitrogen | Cat#10687010 |
| anti-FLAG M2 resin | Sigma | Cat#A2220 |
| Protein A/G resin | Santa Cruz |  |
| polyvinylidene difluoride (PVDF) membrane | GE | Cat#10600022 |
| SuperSignal PicoWest Plus | Thermo/Pierce | Cat#1863099 |
| Experimental Models: Cell Lines |  |  |
| Human: HEK 293 | ATCC |  |
| Human: Jurkat | ATCC |  |
| Human: K562 | ATCC |  |
| Human: KOPTK1 | ATCC |  |
| Human: LOUCY | ATCC |  |
| Human: U937 | ATCC |  |
| Software and Algorithms |  |  |
| Flowjo 10.3 analysis software | FLOWJO, LLC | https://www.flowjo.com/solutions/flowjo |
| Ideas Software | Amnis Corporation | http://www.emdmillipore.com/ |
| ImageLab 5.2.1 | BioRad | https://www.bio-rad.com |
| Imaris | Bitplane Inc |  |
| Other |  |  |
| CytoFLEX benchtop cytometer | Beckman | https://www.beckman.com |
| Leica TCS SP8 confocal imaging system | Leica | https://www.leica-microsystems.com |
| ImageStream MkII | Amnis | https://www.luminexcorp.com |

### Development of a novel multiplexed lentiviral expression vector system

Previously we used multiplexed lentiviral infection with GFP- and RFP-marked viruses to create recombinant leukemia cell lines, in conjunction with fluorescence assisted cell sorting (FACS) (Layer et al., 2016). FACS sorting was laborious and expensive, while the use of GFP and RFP markers limited the number of co-expressed recombinant factors to two (LDB1 and LMO2). Moreover, we observed that initially homogenous FACS-sorted cell lines could inactivate transgene (GFP or RFP) expression over time, consistent with either transgene silencing or competitive advantage/outgrowth of low-expressing clones (JHL and UPD, unpublished). This phenomenon occurred variably amongst different cell lines/types. To circumvent these limitations for the present study, we designed a suite of novel lentiviral vectors. This modular vector family expresses additional fluorescence protein markers that are spectrally distinct, allowing multiplexed co-infection with five or more different viruses. Each vector also encodes a unique antibiotic resistance marker to allow for positive selection of transduced cells. Antibiotic resistance of transduced cells foregoes the need for FACS, and disallows transgene silencing within transduced cell lines; all of which can be proven by antibiotic-enforced consistency of fluorescence marker expression, as monitored by flow cytometry.

### Lentiviral vector construction

We modified a previously described second generation lentiviral vector (Unutmaz et al., 1999). First, an artificial DNA fragment containing the encephalomyocarditis virus internal ribosomal entry site (IRES) sequence, enhanced green fluorescent protein (EGFP) cDNA, and puromycin resistance (PURO) cDNA were assembled *in silico* using publicly available DNA sequences, as follows. A 5’ EcoRI site preceded the IRES sequence, which was immediately followed by a SfiI site flanking the 5’ end of EGFP coding sequence. The initiator methionine codon of EGFP was embedded in the SfiI site. The codon for the last amino acid of EGFP was immediately followed by an NheI site, which immediately preceded the 5’ end of an artificial cDNA encoding human-codon optimized Picornavirus 2A (P2A)-PURO resistance fusion gene. An XhoI site immediately followed the stop codon of the P2A-PURO cassette. This fragment was synthesized as a G Block by Integrated DNA Technologies (IDT, Coralville, Iowa). Synthetic DNA was digested with EcoRI and XhoI and ligated to equivalently digested pBluescript SK (+) (Stratagene). Multiple clonal isolates were subjected to automated DNA sequencing with 5’ M13R and 3’ T7 promoter primers. A single clone perfectly matching the DNA sequence was digested preparatively with EcoRI and XhoI; liberated insert was isolated and ligated to equivalently digested pH110 (Unutmaz et al., 1999). The resultant construct is referred to as pH163-EGFP-PURO. Functionality of pH163 EGFP PURO was first tested for production of virus that could transduce Jurkat cells to EGFP positivity and puromycin resistance (see details below), and the vector backbone was subsequently used as a basis to create additional constructs encoding different combinations of fluorescence markers and antibiotic resistances, as follows. SfiI/NheI fragments corresponding to mCLOVER3, DsREDII, mAPPLE, mSCARLET, EBFPII, mTagBFPII, EYFP, mCITRINE, CERULEAN, mKATE1.3, SMurfBV+, firefly Luciferase, or *S. pyogenes* Cas9 were designed in silico such that non-coding substitutions were made to eliminate any internal NotI, EcoRI, SfiI, NheI, or XhoI sites. Codons were also optimized for human adaptive index on a case-by-case basis, as necessary. mCLOVER3, mSCARLET, mTagBFPII, mKATE1.3, and SMurfBV+ fragments also encoded an amino terminal V5 epitope tag, useful for detection of the recombinant protein in cellular extracts via western blotting. Synthetic G Block DNA was digested with SfiI/NheI and use to replace the equivalent EGFP fragment from H163 EGFP PURO. Insert DNA was verified by automated DNA sequencing, and constructs were tested for functionality according to viral production and transduction/expression within Jurkat cells of the respective fluorescent protein, along with resistance to puromycin.

NheI/XhoI fragments corresponding to P2A-HYGRO, P2A-NEO, P2A-ZEO, and P2A-BLAST were designed in silico according to the above considerations, and synthetic DNAs were used to replace the equivalent P2A-PURO cassette in H163-EGFP-PURO. Individual clonal constructs were validated/tested for ability to produce virus functional for transduction of Jurkat cells to EGFP positivity and resistance to Hygromycin B, G418, Zeocin, or Blasticidin, respectively.

Individual clones conferring the appropriate fluorescent protein expression in combination with PURO selection, or antibiotic resistance companion with EGFP expression, were used to isolate the functionally validated and relevant SfiI/NheI or NheI/XhoI fragment. The isolated functional DNA fragments were used to reconstitute the desired combination of fluorescent marker and antibiotic resistance in the H163 vector backbone, as depicted in FIGURE S1/TABLE X.

### cDNAs and tagged constructs

Subcloning of the 375 amino acid (aa) human LDB1 cDNA was described previously (Layer et al., 2016); wild type cDNA and mutant derivatives were arranged as either 5’ NotI/3’ EcoRI or 5’ BamHI/3’ EcoRI fragments. Vector-embedded epitope tags appended to LDB1 constructs were N-terminal and were either tandem biotin acceptor domain (BAD)/FLAG (MAGGLNDIFEAQKIEWHEGGENLYFQGGDYKDDDDKGGAAASKVRS, FLAG peptide underlined) or HAx1 (MYPYDVPDYAGG). The 158 aa wild type human LMO2 cDNA or mutant derivatives were synthesized as G Blocks with tandem 5’ NotI/BamHI and 3’ EcoRI sites and ligated into NotI/EcoRI digested pBluescript II SK (+). The LMO2 cDNA encoded tandem C-terminal HA (GGMYPYDVPDYA) and SII (GGWSHPQFEK) tags. cDNAs encoding wild type or mutant human 331 aa TAL1, 280 aa LYL1, 361 aa SSBP2, and 388 aa SSBP3 were all synthesized as G Blocks with 5’ NotI/BamHI and 3’ EcoRI sites and ligated into NotI/EcoRI digested pBluescript II SK (+). Sequence encoding N-terminal HAx1 tag (MYPYDVPDYAGG) was located between the 5’ NotI and BamHI sites, and the BamHI site immediately preceded the natural initiator methionine codon. In order to create Lentiviral vectors encoding subunits with BAD/FLAG, HA/SII, or HAx1 tags, clonally-derived NotI/EcoRI fragments encoding BAD/FLAG-LDB1, LMO2-HA/SII, HAx1-TAL1, HAx1-LYL1, HAx1-SSBP2, or HAx1-SSBP3 were transferred from pBluescript II SK (+) vectors into likewise digested H163 vectors. The N-terminal 312 aa Halo tag sequence was PCR amplified from His_6_HaloTag® T7 Vector pH6HTN (Promega) as a 5’ SpeI, 3’ BamHI/EcoRI fragment and ligated into SpeI/EcoRI digested pBluescript II SK (+); the resultant vector was named pHalo-tag-N. Tandem TGA stop codons were located between the BamHI and EcoRI sites. N-terminal HALO fusion constructs were created by ligating clonally-derived BamHI/EcoRI fragments encoding LDB1, LMO2, TAL1, LYL1, SSBP2, or SSBP3 into equivalently digested pHalo-tag-N. In order to create lentiviral vectors encoding N-terminal HALO fusions, NotI/EcoRI fragments were recovered from these pHalo-tag-N vectors and ligated into likewise-digested H163 vectors in order to create H163-Halo-tag-N subunit vectors. All recombinant DNA manipulation and propagation utilized *E. coli* XL1 Blue. All clonal inserts were verified in their entirety by automated DNA sequencing. All mutant derivatives used optimal human codons to encode amino acid substitutions. Maxipreps of lentiviral vector DNA for transfection/virus production were prepared by a modified alkaline lysis/lithium chloride/PEG precipitation protocol in conjunction with extensive phenol/chloroform extraction and ethanol precipitation. Additional details regarding constructs or protocols are available upon request.

### Cell lines, tissue culture, recombinant lentiviruses, transductions, and production of stable cell lines

HEK 293T, Jurkat, K562, U937, KOPT-K1, and LOUCY cells were acquired from the American Type Culture Collection (ATCC). HEK293T cells were cultured in Iscove’s modified Dulbecco’s medium (IMDM)–10% fetal bovine serum (FBS), and other lines were cultured in RPMI 1640–10% FBS, at 37°C in 5% CO_2_. Log-phase HEK 293T cells in 10-cm dishes containing 10 ml medium and 5 × 10^6^ to 8 × 10^6^ cells were transfected by a calcium phosphate–HEPES-buffered saline method with 1 pmol pH163 constructs and 2 pmol pMD-2 for producing pseudotyped lentiviruses. At 12 to 18 h posttransfection, medium was aspirated and replaced with 6 ml fresh medium, which was harvested and replaced at 24 h and 48 h. Media containing viral particles was aliquoted and frozen at −80°C and viral titer was subsequently estimated by serial dilution infection of Jurkat cells. Varying volumes of viral supernatant were mixed with 5 × 10^6^ to 1 × 10^7^ log phase Jurkat cells in a final volume of 10 ml within a T-25 flask (Eppendorf) and subsequently cultured for 72 hours, at which time percentage of fluorescence-positive cells was first roughly determined using an EVOS FL inverted fluorescence microscope (Invitrogen), and then precisely determined using a CytoFLEX benchtop cytometer (Beckman). Microscopy and Cytometry gating parameters were established using parallel culture of non-infected cells as reference. A multiplicity of infection (MOI) of 1 was associated with a fluorescence-positivity of 30% or less. Typical viral titers were 1-2 × 10^6^ infectious particles per milliliter. Jurkat cells infected at an MOI of 1-2 were expanded into a 50 ml culture containing antibiotics to eliminate non-infected cells. Antibiotic regimen and dose varied depending upon the selectable marker encoded by the virus in question and the cell line being transduced; antibiotic concentration kill curves were empirically established for naïve cell lines. As an example, typical antibiotic concentrations for transduced Jurkat cells were puromycin at 2 μg/ml, hygromycin B at 200 μg/ml, G418 at 500 μg/ml, Blasticidin at 10 μg/ml, or Zeocin at 50 μg/ml. After 4-10 days of antibiotic selection cell populations were typically 100% fluorescence positive, at which point they were cryo-preserved in liquid nitrogen using growth media supplemented with 10% DMSO, subjected to iterative rounds of transduction with additional viruses exactly as described above, or used directly for experiments.

### Whole-cell extract, immunoprecipitations, antibodies, and SDS-PAGE/Western blotting

Late-log-phase cultures of ∼7.5 × 10^7^ cells were harvested by centrifugation at 800 × *g* for 10 min, and cell pellets were washed with PBS (phosphate-buffered saline) (2.7 mM KCl, 1.47 mM KH_2_PO_4_, 8.1 mM Na_2_HPO_4_, 137 mM NaCl) and resuspended in 500-1000 μl extraction buffer (20 mM HEPES [pH 7.6], 300 mM NaCl, 20 mM imidazole, 0.1% Triton X-100, 10% glycerol, and protease inhibitor cocktail (Thermo/Pierce)). Cells were disrupted by mild sonication with the microtip of a Branson model 250 sonifier on the low-power setting, and the soluble extract was clarified by centrifugation at 14,000 × *g* for 15 min. Extract protein content was typically 5 to 10 μg/μl. A portion was mixed with an equal volume of 2× SDS sample buffer and briefly heated to 75°C. For immunoprecipitations (IP), 100 μl of soluble extract was supplemented with an additional 100 μl of extraction buffer also containing 5 μl anti-FLAG M2 resin (catalog number A2220; Sigma) or 5 μl of Protein A/G resin (Santa Cruz) along with 1-2 micrograms of anti-LMO2 IgG, then rocked at 4°C for 3 to 4 h. Immune complexes were isolated by centrifugation, washed 3 times with 200 μl of extraction buffer, and eluted by heating with 100 μl SDS sample buffer. Samples were stored at −80°C and briefly heated again at 75°C just prior to loading onto handcast discontinuous SDS-PAGE gels with a 4% acrylamide stacking gel and a 4-to-15% linear gradient resolving gel (37.5%/1.0% [wt/vol] acrylamide-bisacrylamide), run at 15 V/cm for 90-105 min. Gels were transferred onto a 0.2-μm polyvinylidene difluoride (PVDF) membrane (catalog number 10600022; GE) at 50 V for 2.5 h; filters were blocked in PBS–2% non fat dry milk (NFDM, Marsh FoodClub) and incubated with antibodies in blocking buffer overnight at 4°C.

The following antibodies for Western blotting were used according to the manufacturer’s specifications: mouse monoclonal anti LDB1 IgG (catalog number sc-376030x; Santa Cruz) (detected with a goat anti mouse IgG Fc-horseradish peroxidase (HRP) conjugate, catalog number 31439; Thermo/Pierce), anti FLAG-HRP conjugate (catalog number A8592; Sigma), anti HA-HRP conjugate (catalog number 12013819001; Roche), anti V5-HRP conjugate (to detect mSCARLET and other V5 tagged fluorescent proteins, catalog number 46-0708, Invitrogen), rabbit polyclonal anti TAL1 IgG (catalog number A305-300A, Bethyl), (detected with a goat anti rabbit IgG-HRP conjugate [catalog number 211-032-171; Jackson ImmunoResearch]), mouse monoclonal anti SSBP2 IgG (catalog number sc-166687, Santa Cruz), mouse monoclonal anti HALO IgG (catalog number G921A, Promega), mouse monoclonal anti GFP IgG (catalog number 11814460001; Roche), rabbit polyclonal anti tubulin IgG (catalog number SC-9104; Santa Cruz). The high-affinity/sensitivity/specificity mouse anti valosin-containing protein (anti VCP) antibody (catalog number ab11433; Abcam) was used for multiplex Western blotting as a loading control. The 1A93B11 mouse anti LMO2 IgG was described previously (Layer et al., 2016).

Western blots were developed with enhanced chemiluminescense (ECL) detection (SuperSignal Pico West Plus, catalog number 1863099, Thermo/Pierce). All images were obtained within the linear signal detection range using a ChemiDoc Touch imaging system (BioRad). Images were analyzed using ImageLab Software version 5.2.1 (BioRad) and exported to Adobe Photoshop and Illustrator for figure assembly.

### HaloLife assay: live cell pulse chase analysis

1.25 10^5^ cells were collected from log-phase cultures by centrifugation at 1,200 *x g* for 1 min. The culture media was removed, and cells were resuspended with 125 µL RPMI containing 10% FBS and HaloTag Ligand R110 (Promega Ca.) at a final concentration of 100nM, per the company’s instructions. The resuspended cells were then incubated for 90 min at 37 °C in 5% CO2. After 90 min the cells were centrifuged at 12,000 x g for 1 min and washed with PBS (2.7 mM KCl, 1.47 mM KH2PO4, 8.1 mM Na2HPO4, 137 NaCl) containing 0.1% BSA (bovine serum albumin) a total of 3 times to remove excess HaloTag Ligand R110. Cells were resuspended in 600 µL RPMI containing 10% FBS, and 4, 150 µL aliquots were transferred to a 96-well round-bottom plate (TPP). 10,000 events were then immediately analyzed from 1 of the 4 150 µL aliquots using a CytoFLEX benchtop cytometer (Beckman). All subsequent chase time points were collected using this initial analysis as a reference. Between flow cytometry analyses, the 96-well plate containing the HaloTag Ligand R110 labeled cells were placed in an incubator at 37°C with 5% CO2 until the next collection point. Flow cytometry analyses were collected 3, 4, and 5 hours after T0 for all cells, with the exception those containing Halo-tagged LDB1 and LYL1 due to their significantly different observed half-lives. For cells containing Halo-tagged LDB1, flow cytometry events were recorded at 6, 12, and 24 hours after T0, and analyses were recorded 1, 2, and 3 hours after the initial time point for cells containing Halo-tagged LYL1. Replicate experiments were done on consecutive days.

### Pulse-chase FCS file analysis

All FCS files were analyzed using Flowjo 10.3 analysis software (FLOWJO, LLC, OR). To identify cells that were co-expressing EBFPII and/or mScarlet in conjunction with Halo-tagged proteins, non-transduced unstained Jurkat cells were used to establish a gating sequence. Their physical dimensions were grouped on an FSC-A/FSC-H plot to determine the total number of lymphocytes within the event population. A gate was then established on an FSC-A/SSC-A plot to select for live cells within the total lymphocyte population. The resulting population was then gated as a negative control for both fluorescence markers on a PB450-A (EBFPII)/FITC-A (HaloTag R110) plot. This gating sequence was then applied to all FCS files within the same experiment.

### Half-Life Calculations

Log-linear regression curves were calculated from flow cytometry analysis data to calculate Halo-tagged protein half-lives. PB450-A (EBFPII) and FITC-A (HaloTag R110 Ligand) double positive events were calculated as a percentage of the parent population for all time points collected. Replicate data for each time point was averaged, and then normalized to the initial time point. The natural log was calculated for each of the averages, and the resulting values were represented over time on a 2-dimensional scatter plot. A trend line was calculated, and the resulting slope was used to determine Halo-tagged protein half-lives.

### Statistical Analysis

The standard error of the mean (SEM) was calculated for individual time points in each Halo-tagged protein experiment using Microsoft Excel. SEM values were then applied to their corresponding time points within the log-linear regression curves used to determine Halo-tagged protein half-lives. Results from replicate experiments were used to calculate the standard deviation, which was then divided by the square root of the number of replicates to determine the SEM. The SEM for Halo-tagged protein half-lives values were also calculated using the same formula. Half-life values were analyzed from at least 3 experiments, as previously described, and then used to calculate the SEM.

### ImageStream

1.25 10^5^ cells were collected from log-phase cultures by centrifugation at 1,200 x g for 1 min. The culture media was removed, and cells were resuspended with 125 µL RPMI containing 10% FBS and HaloTag Ligand R110 (Promega Ca.) at a final concentration of 100nM, per the company’s instructions. The resuspended cells were then incubated for 90 min at 37°C in 5% CO2. After 90 min the cells were centrifuged at 12,000 x g for 1 min and washed with PBS (2.7 mM KCl, 1.47 mM KH2PO4, 8.1 mM Na2HPO4, 137 NaCl) containing 0.1% BSA (bovine serum albumin) a total of 3 times to remove excess HaloTag Ligand R110. The cells were then resuspended in 1mL PBS, and stained with SYTO 17 red fluorescent nucleic acid stain (Invitrogen) at a final concentration of 10 nM for 10 min, per manufacturer’s instructions. The cells were washed once more, and resuspended with 200 µL PBS before being analyzed using ImageStream®^X^ Mark II Imaging Flow Cytometer (MilliporeSigma). Data analysis was done using the IDEAS 6.2’s (Millipore) nuclear localization analysis Wizard.

### Confocal Imaging

1.25×10^5^ cells were collected from log-phase cultures by centrifugation at 1,200 x g for 1 min. The culture media was removed, and cells were resuspended with 125 µL RPMI containing 10% FBS and HaloTag Ligand R110 (Promega Ca.) at a final concentration of 100nM, then incubated for 90 min at 37°C in 5% CO2. After 90 min the HaloTag Ligand R110 labeled cells were centrifuged at 1,200 x g for 1 min and washed with PBS (2.7 mM KCl, 1.47 mM KH2PO4, 8.1 mM Na2HPO4, 137 NaCl) containing 0.1% BSA (bovine serum albumin) a total of 3 times to remove excess ligand. The washed cells were then resuspended in 1mL of PBS and stained with SYTO 17 red fluorescent nucleic acid stain (Molecular Probes, Inc. OR) according to the manufacturer’s protocol. After the incubation period, the cells were centrifuged at 1,200 x g for 1 min and washed once with PBS. Once resuspended in 300 µL of PBS, cells were transferred to a 12 mm glass base dish and imaged with a Leica TCS SP8 confocal imaging system (Leica Microsystems Inc, IL) using an HC PL APO 40x/1.3 oil CS2 objective. Digital images were rendered, and signal intensities were analyzed using Imaris visualization and analysis software (Bitplane Inc. MA). Cellular localization of HaloTaged proteins was determined by calculating the ratio of mean HaloTag signal intensities within the nucleus versus the cytosol. The nuclear area was established using the SYTO 17 red fluorescent nucleic acid stain, and the cytoplasmic region was determined using the diffuse EBFPII signal expressed by our lentiviral vectors.

## References

Bonifacino, J.S., Cosson, P., and Klausner, R.D. (1990). Colocalized transmembrane determinants for ER degradation and subunit assembly explain the intracellular fate of TCR chains. Cell 63, 503–513.

Breitschopf, K., Bengal, E., Ziv, T., Admon, A., and Ciechanover, A. (1998). A novel site for ubiquitination: the N-terminal residue, and not internal lysines of MyoD, is essential for conjugation and degradation of the protein. The EMBO Journal 17, 5964.

Cai, G., Imasaki, T., Yamada, K., Cardelli, F., Takagi, Y., and Asturias, F.J. (2010). Mediator head module structure and functional interactions. Nat Struct Mol Biol 17, 273–279.

Davé, U., Akagi, K., Tripathi, R., and Cleveland, S. (2009). Murine leukemias with retroviral insertions at Lmo2 are predictive of the leukemias induced in SCID-X1 patients following retroviral gene therapy. PLoS genetics.

Davé, U.P., Jenkins, N.A., and Copeland, N.G. (2004). Gene therapy insertional mutagenesis insights. Science 303, 333.

Deng, W., Lee, J., Wang, H., Miller, J., Reik, A., Gregory, P.D., Dean, A., and Blobel, G.A. (2012). Controlling long-range genomic interactions at a native locus by targeted tethering of a looping factor. Cell 149, 1233–1244.

Deshaies, R.J. (2015). Protein degradation: prime time for PROTACs. Nature chemical biology 11, 634.

Dong, W.F., Xu, Y., Hu, Q.L., Munroe, D., Minowada, J., Housman, D.E., and Minden, M.D. (1995). Molecular characterization of a chromosome translocation breakpoint t(11;14)(p13;q11) from the cell line KOPT-K1. Leukemia 9, 1812–1817.

El Omari, K., Hoosdally, S.J., Tuladhar, K., Karia, D., Vyas, P., Patient, R., Porcher, C., and Mancini, E.J. (2011). Structure of the leukemia oncogene LMO2: implications for the assembly of a hematopoietic transcription factor complex. Blood 117, 2146–2156.

Emanuele, Michael J., Elia, Andrew E.H., Xu, Q., Thoma, Claudio R., Izhar, L., Leng, Y., Guo, A., Chen, Y.-N., Rush, J., Hsu, Paul W.-C., et al. (2011). Global Identification of Modular Cullin-RING Ligase Substrates. Cell 147, 459–474.

Ferrando, A.A., Neuberg, D.S., Staunton, J., Loh, M.L., Huard, C., Raimondi, S.C., Behm, F.G., Pui, C.H., Downing, J.R., Gilliland, D.G., et al. (2002). Gene expression signatures define novel oncogenic pathways in T cell acute lymphoblastic leukemia. Cancer Cell 1, 75–87.

Gao, X.-D., Tachikawa, H., Sato, T., Jigami, Y., and Dean, N. (2005). Alg14 Recruits Alg13 to the Cytoplasmic Face of the Endoplasmic Reticulum to Form a Novel Bipartite UDP-N-acetylglucosamine Transferase Required for the Second Step of N-Linked Glycosylation. Journal of Biological Chemistry 280, 36254–36262-36262.

Greer, J.P. (2019). Wintrobe’s clinical hematology, Fourteenth edition. edn (Philadelphia, PA: Wolters Kluwer).

Güngör, C., Taniguchi-Ishigaki, N., Ma, H., Drung, A., Tursun, B., Ostendorff, H.P., Bossenz, M., Becker, C.G., Becker, T., and Bach, I. (2007). Proteasomal selection of multiprotein complexes recruited by LIM homeodomain transcription factors. Proceedings of the National Academy of Sciences 104, 15000–15005.

Hewitt, K., Johnson, K., Gao, X., Keles, S., and Bresnick, E. (2016). The hematopoietic stem and progenitor cell cistrome: GATA factor-dependent cis-regulatory mechanisms. In Current topics in developmental biology (Elsevier), pp. 45–76.

Hewitt, Kyle J., Kim, Duk H., Devadas, P., Prathibha, R., Zuo, C., Sanalkumar, R., Johnson, Kirby D., Kang, Y.-A., Kim, J.-S., Dewey, Colin N., et al. (2015). Hematopoietic Signaling Mechanism Revealed from a Stem/Progenitor Cell Cistrome. Molecular Cell 59, 62–74.

Howard, P.W., Jue, S.F., Ransom, D.G., and Maurer, R.A. (2010). Regulation of LIM-domain-binding 1 protein expression by ubiquitination of Lys 134. The Biochemical journal 429, 127–136.

Imasaki, T., Calero, G., Cai, G., Tsai, K.L., Yamada, K., Cardelli, F., Erdjument-Bromage, H., Tempst, P., Berger, I., Kornberg, G.L., et al. (2011). Architecture of the Mediator head module. Nature 475, 240–243.

Jagannathan-Bogdan, M., and Zon, L.I. (2013). Hematopoiesis. Development 140, 2463–2467.

Kim, W., Bennett, Eric J., Huttlin, Edward L., Guo, A., Li, J., Possemato, A., Sowa, Mathew E., Rad, R., Rush, J., Comb, Michael J., et al. (2011). Systematic and Quantitative Assessment of the Ubiquitin-Modified Proteome. Molecular Cell 44, 325–340.

Komander, D., and Rape, M. (2012). The ubiquitin code. Annual review of biochemistry 81, 203–229.

Krivega, I., Dale, R.K., and Dean, A. (2014a). Role of LDB1 in the transition from chromatin looping to transcription activation. Genes & development 28, 1278–1290.

Krivega, I., Dale, R.K., and Dean, A. (2014b). Role of LDB1 in the transition from chromatin looping to transcription activation. Genes Dev 28, 1278–1290.

Lai, A.C., and Crews, C.M. (2017). Induced protein degradation: an emerging drug discovery paradigm. Nature reviews Drug discovery 16, 101.

Layer, J.H., Alford, C.E., McDonald, W.H., and Dave, U.P. (2016). LMO2 Oncoprotein Stability in T-Cell Leukemia Requires Direct LDB1 Binding. Mol Cell Biol 36, 488–506.

Li, L., Jothi, R., Cui, K., Lee, J.Y., Cohen, T., Gorivodsky, M., Tzchori, I., Zhao, Y., Hayes, S.M., Bresnick, E.H., et al. (2011). Nuclear adaptor Ldb1 regulates a transcriptional program essential for the maintenance of hematopoietic stem cells. Nat Immunol 12, 129–136.

Liu, G., and Dean, A. (2019). Enhancer long-range contacts: The multi-adaptor protein LDB1 is the tie that binds. Biochimica et Biophysica Acta (BBA)-Gene Regulatory Mechanisms.

Los, G.V., Encell, L.P., McDougall, M.G., Hartzell, D.D., Karassina, N., Zimprich, C., Wood, M.G., Learish, R., Ohana, R.F., and Urh, M. (2008). HaloTag: a novel protein labeling technology for cell imaging and protein analysis. ACS chemical biology 3, 373–382.

Lurie, L.J., Boyer, M.E., Grass, J.A., and Bresnick, E.H. (2008). Differential GATA factor stabilities: implications for chromatin occupancy by structurally similar transcription factors. Biochemistry 47, 859–869.

Mashtalir, N., D’Avino, A.R., Michel, B.C., Luo, J., Pan, J., Otto, J.E., Zullow, H.J., McKenzie, Z.M., Kubiak, R.L., and Pierre, R.S. (2018). Modular organization and assembly of SWI/SNF family chromatin remodeling complexes. Cell 175, 1272–1288. e1220.

McCormack, M.P., and Rabbitts, T.H. (2004). Activation of the T-cell oncogene LMO2 after gene therapy for X-linked severe combined immunodeficiency. N Engl J Med 350, 913–922.

McCormack, M.P., Shields, B.J., Jackson, J.T., Nasa, C., Shi, W., Slater, N.J., Tremblay, C.S., Rabbitts, T.H., and Curtis, D.J. (2013). Requirement for Lyl1 in a model of Lmo2-driven early T-cell precursor ALL. Blood 122, 2093–2103.

Meier, N., Krpic, S., Rodriguez, P., Strouboulis, J., Monti, M., Krijgsveld, J., Gering, M., Patient, R., Hostert, A., and Grosveld, F. (2006). Novel binding partners of Ldb1 are required for haematopoietic development. Development (Cambridge, England) 133, 4913–4923.

Mevissen, T.E., Hospenthal, M.K., Geurink, P.P., Elliott, P.R., Akutsu, M., Arnaudo, N., Ekkebus, R., Kulathu, Y., Wauer, T., and El Oualid, F. (2013). OTU deubiquitinases reveal mechanisms of linkage specificity and enable ubiquitin chain restriction analysis. Cell 154, 169–184.

Morrow, M.E., Morgan, M.T., Clerici, M., Growkova, K., Yan, M., Komander, D., Sixma, T.K., Simicek, M., and Wolberger, C. (2018). Active site alanine mutations convert deubiquitinases into high-affinity ubiquitin-binding proteins. EMBO reports 19, e45680.

Murre, C. (2019). Helix–loop–helix proteins and the advent of cellular diversity: 30 years of discovery. Genes & development 33, 6–25.

Nam, C.H., and Rabbitts, T.H. (2006). The role of LMO2 in development and in T cell leukemia after chromosomal translocation or retroviral insertion. Mol Ther 13, 15–25.

Ono, Y., Fukuhara, N., and Yoshie, O. (1998). TAL1 and LIM-only proteins synergistically induce retinaldehyde dehydrogenase 2 expression in T-cell acute lymphoblastic leukemia by acting as cofactors for GATA3. Molecular and cellular biology 18, 6939–6950.

Orkin, S.H., and Zon, L.I. (2008). SnapShot: hematopoiesis. Cell 132, 712–712.

Ostendorff, H.P., Peirano, R.I., Peters, M.A., Schluter, A., Bossenz, M., Scheffner, M., and Bach, I. (2002). Ubiquitination-dependent cofactor exchange on LIM homeodomain transcription factors. Nature 416, 99–103.

Raetz, E.A., and Teachey, D.T. (2016). T-cell acute lymphoblastic leukemia. ASH Education Program Book 2016, 580–588.

Schlaeger, T.M., Schuh, A., Flitter, S., Fisher, A., Mikkola, H., Orkin, S.H., Vyas, P., and Porcher, C. (2004). Decoding hematopoietic specificity in the helix-loop-helix domain of the transcription factor SCL/Tal-1. Mol Cell Biol 24, 7491–7502.

Smith, S., Tripathi, R., Goodings, C., Cleveland, S., Mathias, E., Hardaway, J.A., Elliott, N., Yi, Y., Chen, X., Downing, J., et al. (2014). LIM domain only-2 (LMO2) induces T-cell leukemia by two distinct pathways. PLoS ONE 9, e85883.

Soler, E., Andrieu-Soler, C., de Boer, E., Bryne, J.C., Thongjuea, S., Stadhouders, R., Palstra, R.-J., Stevens, M., Kockx, C., and van IJcken, W. (2010). The genome-wide dynamics of the binding of Ldb1 complexes during erythroid differentiation. Genes & development 24, 277–289.

Song, S.-H., Hou, C., and Dean, A. (2007). A positive role for NLI/Ldb1 in long-range β-globin locus control region function. Molecular cell 28, 810–822.

Sun, X.-J., Wang, Z., Wang, L., Jiang, Y., Kost, N., Soong, T.D., Chen, W.-Y., Tang, Z., Nakadai, T., Elemento, O., et al. (2013). A stable transcription factor complex nucleated by oligomeric AML1-ETO controls leukaemogenesis. Nature, 1–6.

Trausch-Azar, J.S., Lingbeck, J., Ciechanover, A., and Schwartz, A.L. (2004). Ubiquitin-Proteasome-mediated degradation of Id1 is modulated by MyoD. Journal of Biological Chemistry 279, 32614–32619.

Unutmaz, D., KewalRamani, V.N., Marmon, S., and Littman, D.R. (1999). Cytokine signals are sufficient for HIV-1 infection of resting human T lymphocytes. 10.1084/jem20050075 189, 1735–1746.

Vilimas, T., Mascarenhas, J., Palomero, T., Mandal, M., Buonamici, S., Meng, F., Thompson, B., Spaulding, C., Macaroun, S., and Alegre, M.-L. (2007). Targeting the NF-κB signaling pathway in Notch1-induced T-cell leukemia. Nature medicine 13, 70.

Wadman, I.A., Osada, H., Grutz, G.G., Agulnick, A.D., Westphal, H., Forster, A., and Rabbitts, T.H. (1997). The LIM-only protein Lmo2 is a bridging molecule assembling an erythroid, DNA-binding complex which includes the TAL1, E47, GATA-1 and Ldb1/NLI proteins. Embo J 16, 3145–3157.

Wang, T., Yu, H., Hughes, N.W., Liu, B., Kendirli, A., Klein, K., Chen, W.W., Lander, E.S., and Sabatini, D.M. (2017). Gene essentiality profiling reveals gene networks and synthetic lethal interactions with oncogenic Ras. Cell 168, 890–903. e815.

Wilson, N.K., Foster, S.D., Wang, X., Knezevic, K., Schütte, J., Kaimakis, P., Chilarska, P.M., Kinston, S., Ouwehand, W.H., and Dzierzak, E. (2010). Combinatorial transcriptional control in blood stem/progenitor cells: genome-wide analysis of ten major transcriptional regulators. Cell stem cell 7, 532–544.

Wright, K.J., Marr, M.T., and Tjian, R. (2006). TAF4 nucleates a core subcomplex of TFIID and mediates activated transcription from a TATA-less promoter. Proceedings of the National Academy of Sciences 103, 12347–12352.

Xu, Z., Huang, S., Chang, L.S., Agulnick, A.D., and Brandt, S.J. (2003). Identification of a TAL1 target gene reveals a positive role for the LIM domain-binding protein Ldb1 in erythroid gene expression and differentiation. Mol Cell Biol 23, 7585–7599.

Xu, Z., Meng, X., Cai, Y., Liang, H., Nagarajan, L., and Brandt, S.J. (2007). Single-stranded DNA-binding proteins regulate the abundance of LIM domain and LIM domain-binding proteins. Genes Dev 21, 942–955.

