## Supplemental Figures for "Live cell kinetic analysis of the LMO2/LDB1 leukemogenic protein complex reveals a hierarchy of turnover with implications for complex assembly"

**SUPPLEMENTAL INFORMATION**

**Figure S1 refers to Figure 1 and STAR methods section. Lentiviral constructs that allow multiplexing.**

(A) Schematic shows the structure of the lentiviral vectors used in this study. See STAR methods for details on construction of this vector from its backbone. Basically, we have made multiple vectors with combinations of fluorescent protein tags and drug selection. The excitation, emission, and brightness of each fluorescent protein is shown.

**Figure S2 refers to Figure 2. Co-IP of LMO2 and LDB1 and mutant proteins shows lysineless LMO2 is deficient in binding to LDB1.**

(A) Clustal alignment of LIM domain Only paralogs, LMO1, LMO2, and LMO4. Arrows denote lysine residues throughout LMO2 protein.

(B) Immunoblot with various antibodies of transduced Jurkat cell lysate. Expression control (GFP, mScarlet) and loading controls (VCP) are included. LDB1 WT and mutant proteins, LDB1ΔLID, LDB1(I322A), LMO2 WT, and LMO2 K(0), a lysineless version of LMO2 with arginine substituted for each lysine.

(C) Immunoprecipitation from same Jurkat cell lysate with anti-FLAG to pull down FLAG-LDB1. IgG capture is shown with blots for Fc and light chain.

(D) Immunoprecipitation from same Jurkat cell lysate with anti-LMO2 antibody to pull down LMO2.

**Figure S3 refers to Figure 4. SSBP2 and SSBP3 half-lives and immunoblots.**

(A) Immunoblot of transduced Jurkat cells with various antibodies. SSBP2 and SSBP3 were N-terminally tagged with Halo and co-expressed with LDB1 WT or various mutant LDB1 proteins, LDB1ΔLCCD is deficient in binding to SSBP proteins; K134R and K365R are arginine substitutions that can stabilize LDB1 from degradation. Expression control (EBFPII) and loading controls (tubulin and VCP) are included.

(B) HaloLife assay was performed in Jurkat cells for Halo-SSBP2 with $t_{1/2}$ shown.

(C) HaloLife assay was performed in Jurkat cells for Halo-SSBP3 with $t_{1/2}$ shown. Table in inset shows the lysines within SSBP proteins are located within the LUFS domain responsible for binding to LDB1 protein.

**Figure S4 refers to Figure 7. Full table of DUB screen using HaloLife assay.** Top left panel shows schematic of experiment for screening DUBs in the HaloLife assay. The effect of DUB shRNA knockdown was analyzed for each Halo-tagged subunit protein. In the following tables, DUBs are grouped in families. First column shows whether they are expressed in Jurkat cells followed by their effect upon the stability of Halo-tagged proteins. Yellow denotes 2 out of 3 criteria met. Red shows 3 out of 3 criteria met.

**Figure S5 refers to discussion section. Confocal imaging of Halo-tagged proteins.** Confocal microscopy images of Jurkat cells showing localization of Halo-tagged proteins. Cells were transduced with our multiplexed lentiviral expression vectors to express HALO-tagged LMO2, LDB1, Tal1, Lyl1, SSBP2, and SSBP3, respectively. Non-transduced Jurkat cells were imaged as a negative control, shown in first row. Cellular localization of Halo-tagged proteins was determined by calculating the ratio of mean HaloTag R110 signal intensities within the nucleus versus the cytosol. Nuclear and cytosolic regions were established from SYTO Red 17 and EBFPII signals, respectively.

FIGURE S1

Justin H. Layer and Utpal P. Dave
Multiplex Lentiviral Expression Vector System
2018 Version

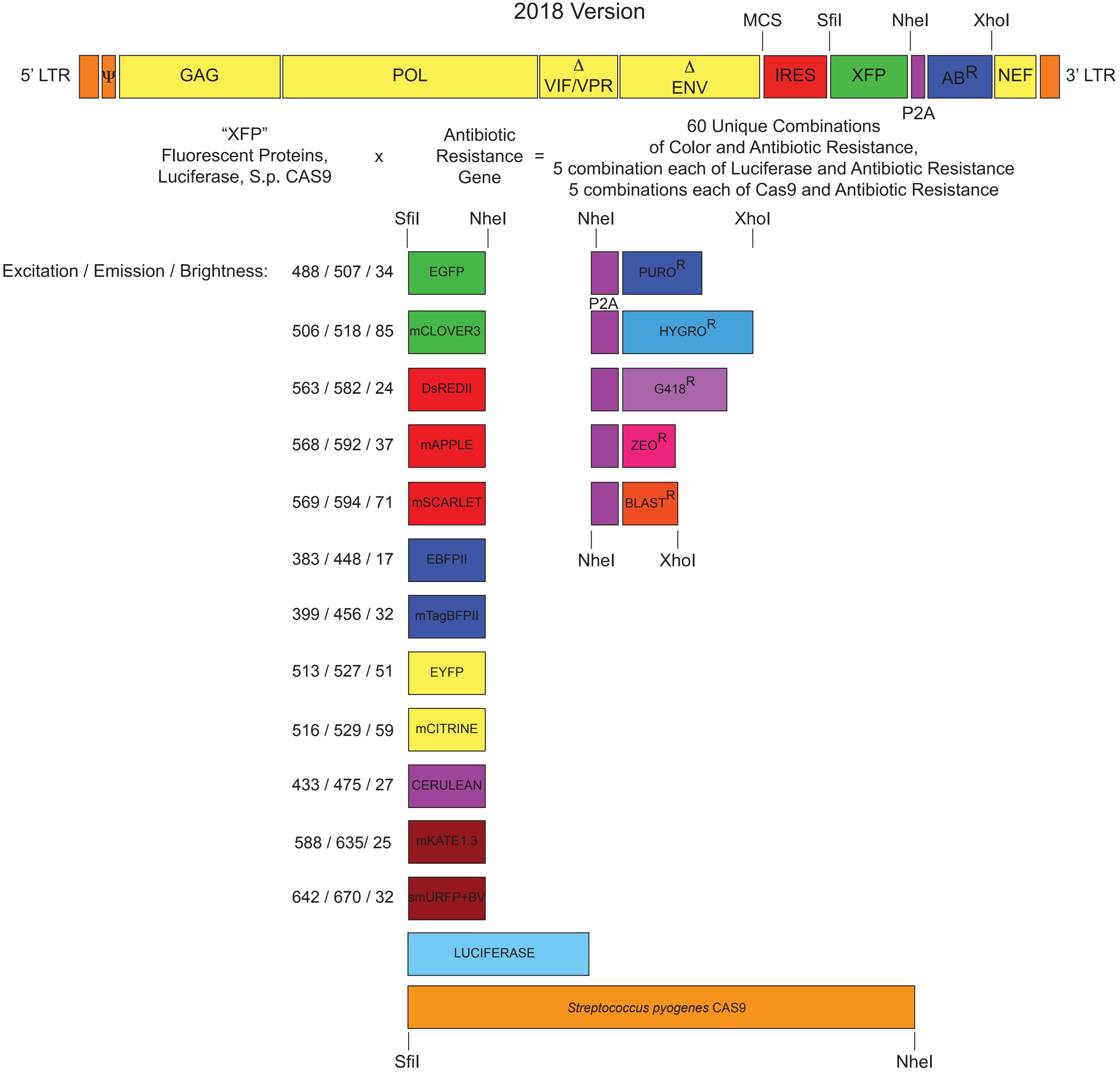

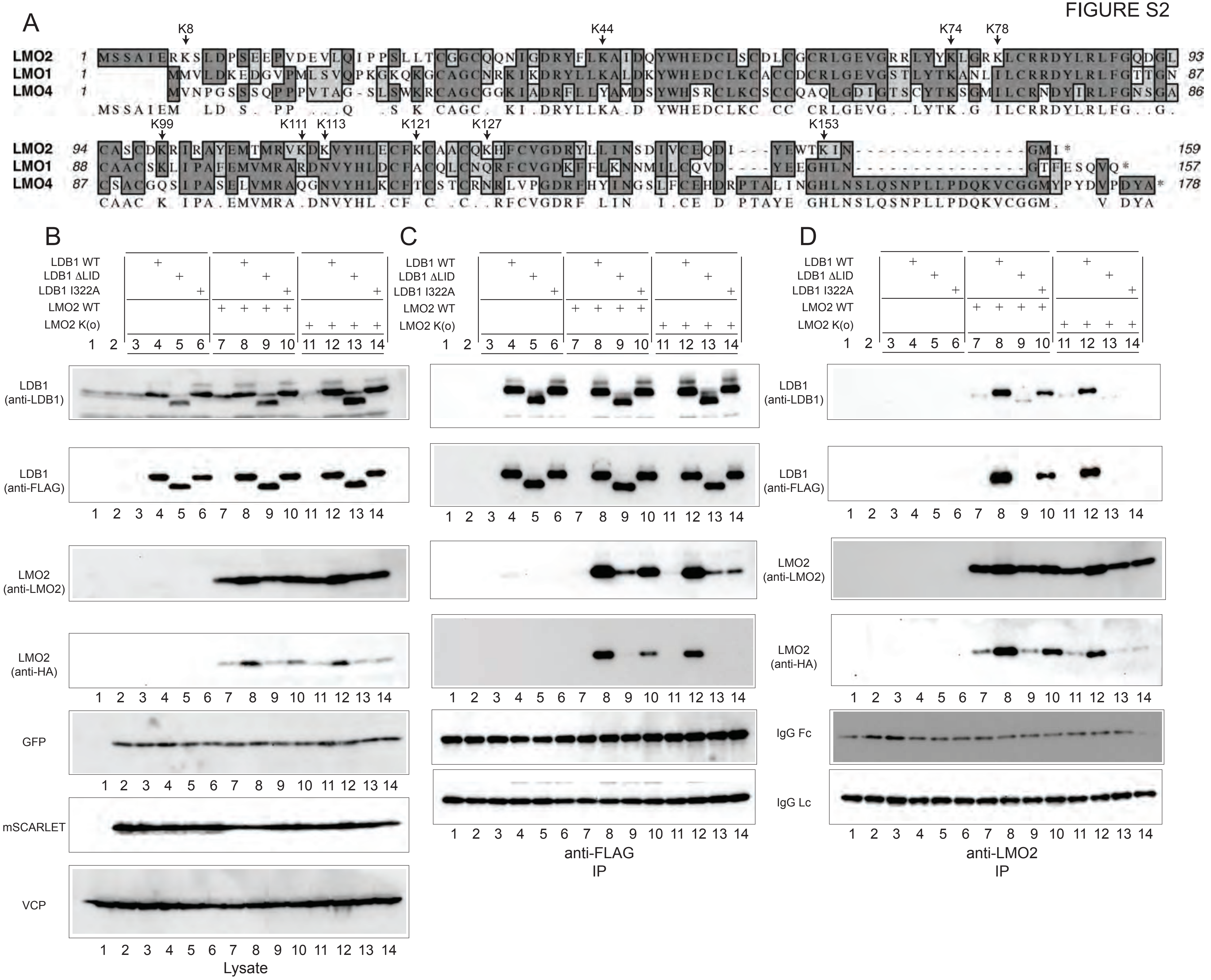
A
K8
K44
K74
K78
LMO2
LMO1
LMO4
K99
K111
K113
K121
K127
K153
B
C
D
LDB1 WT
LDB1 ΔLID
LDB1 I322A
LMO2 WT
LMO2 K(o)
LDB1 (anti-LDB1)
LDB1 (anti-FLAG)
LMO2 (anti-LMO2)
LMO2 (anti-HA)
GFP
mSCARLET
VCP
Lysate
IgG Fc
IgG Lc
anti-FLAG IP
anti-LMO2 IP

FIGURE S3

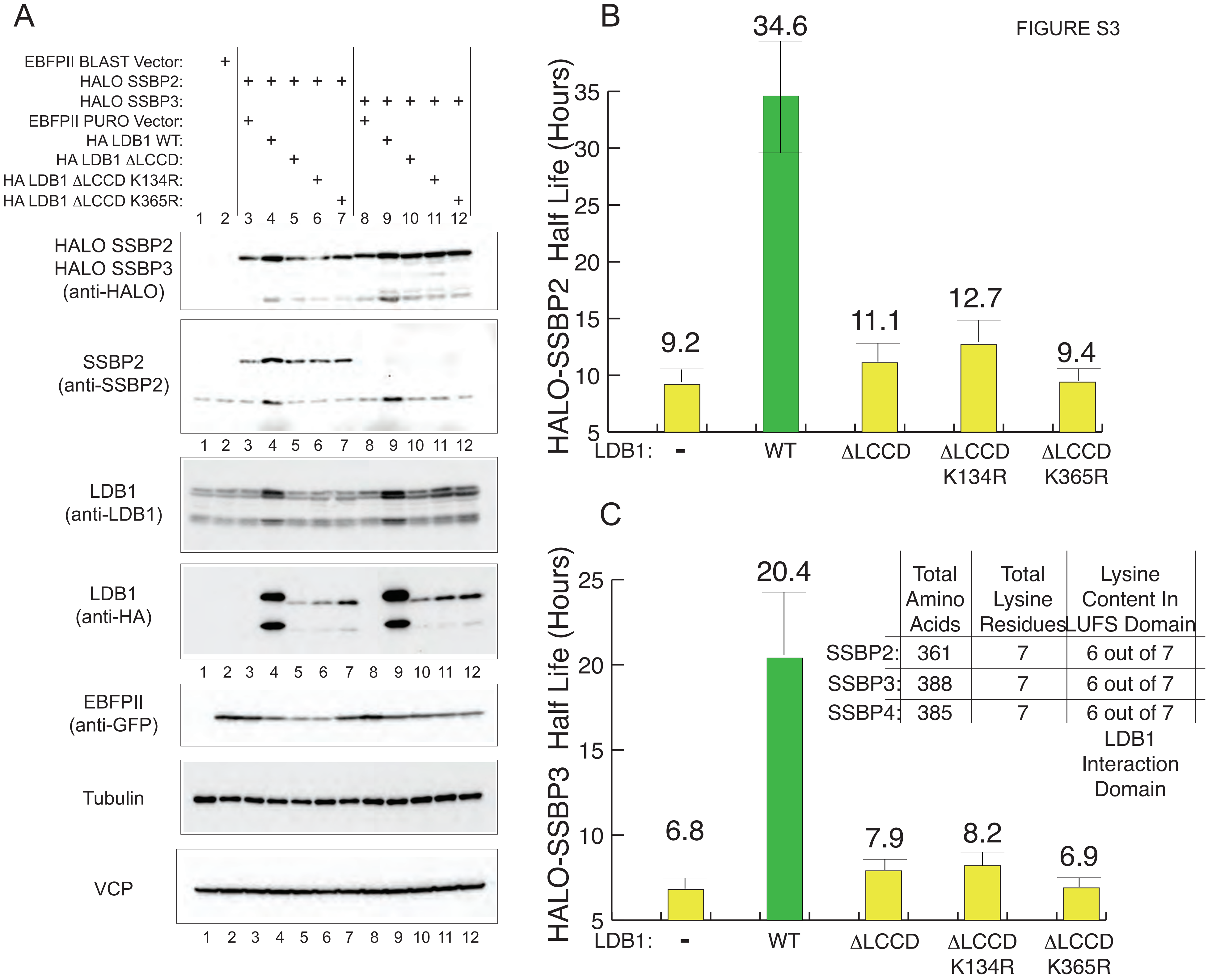
A
EBFPII BLAST Vector:
HALO SSBP2:
HALO SSBP3:
EBFPII PURO Vector:
HA LDB1 WT:
HA LDB1 ΔLCCD:
HA LDB1 ΔLCCD K134R:
HA LDB1 ΔLCCD K365R:
1 2 3 4 5 6 7 8 9 10 11 12
HALO SSBP2
HALO SSBP3
(anti-HALO)
SSBP2
(anti-SSBP2)
LDB1
(anti-LDB1)
LDB1
(anti-HA)
EBFPII
(anti-GFP)
Tubulin
VCP
B
HALO-SSBP2 Half Life (Hours)
34.6
9.2
11.1
12.7
9.4
LDB1: - WT ΔLCCD ΔLCCD K134R ΔLCCD K365R
C
HALO-SSBP3 Half Life (Hours)
20.4
6.8
7.9
8.2
6.9
LDB1: - WT ΔLCCD ΔLCCD K134R ΔLCCD K365R
Total Amino Acids
Total Lysine Residues
Lysine Content In LUFS Domain
SSBP2: 361 7 6 out of 7
SSBP3: 388 7 6 out of 7
SSBP4: 385 7 6 out of 7
LDB1 Interaction Domain

### FIGURE S4

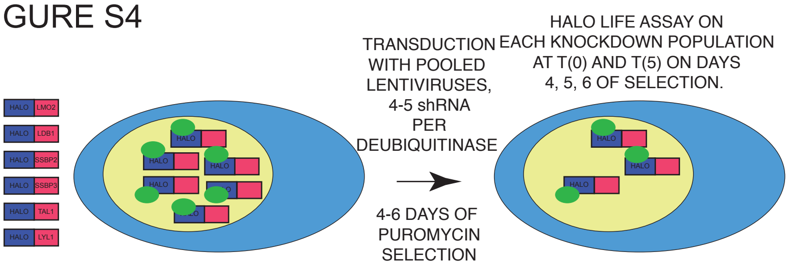


POSITIVE HITS SCORED BY:

I. REDUCED % OF T(0) DUB shRNA TO T(0) SCRAMBLED shRNA

II. REDUCED T(0) HALO SIGNAL

III. REDUCED % OF T(0) at T(5)

FULFILLS AT LEAST 2 CRITERIA

FULFILLS ALL 3 CRITERIA

MISSING shRNA (14):
USP19
USP24
USP27
USP35
USP45
FAM63A
OTULIN
OTU1
OTUD1
OTUD3
OTUD4
AT7L3
JOSD2
ZUFSP

#### OTU FAMILY (11 Tested)

| | Present in JURKAT? | ΣΛΟΩ ΓΡΟΩΤΗ? | HALO LMO2 | HALO LDB1 | HALO SSBP2 | HALO SSBP3 | HALO TAL1 | HALO LYL1 |
|---|---|---|---|---|---|---|---|---|
| OTUB1 | NO | NO | | | | | | |
| OTUB2 | YES | NO | | | YES | | | |
| OTUD5 | YES | NO | | | | | | |
| OTUD6A | ? | NO | | | | YES | | |
| OTUD6B | YES | YES | | | | | YES | |
| OTUD7A | YES | YES | | | | | | |
| OTUD7B | YES | NO | YES | YES | YES | YES | YES | YES |
| ALG13 | YES | SEVERE | YES | YES | YES | YES | YES | YES |
| TNFAIP3 | ? | SEVERE | | | | | | |
| VCPIP1 | YES | NO | | | | | | YES |
| ZRANB1 | ? | NO | | | YES | | | YES |
| | | | HALO LMO2 | HALO LDB1 | HALO SSBP2 | HALO SSBP3 | HALO TAL1 | HALO LYL1 |

#### MINDY FAMILY (5 Tested)

| | Present in JURKAT? | ΣΛΟΩ ΓΡΟΩΤΗ? | HALO LMO2 | HALO LDB1 | HALO SSBP2 | HALO SSBP3 | HALO TAL1 | HALO LYL1 |
|---|---|---|---|---|---|---|---|---|
| FAM63B | YES | YES | | | | | YES | YES |
| FAM105A | ? | NO | | | | | | |
| FAM105B | ? | NO | | | YES | YES | | YES |
| FAM188A | YES | YES | | | | | YES | |
| FAM188B | YES | YES | | | | | | YES |

#### UCHL FAMILY (3 Tested)

| | Present in JURKAT? | ΣΛΟΩ ΓΡΟΩΤΗ? | HALO LMO2 | HALO LDB1 | HALO SSBP2 | HALO SSBP3 | HALO TAL1 | HALO LYL1 |
|---|---|---|---|---|---|---|---|---|
| UCHL1 | NO | YES | | YES | YES | YES | | |
| UCHL3 | YES | NO | | | YES | | | |
| UCHL5 | YES | NO | | | | | | |
| BAP1 | YES | N.D. | | | | | | |

#### MJD FAMILY (1 Tested)

| | Present in JURKAT? | ΣΛΟΩ ΓΡΟΩΤΗ? | HALO LMO2 | HALO LDB1 | HALO SSBP2 | HALO SSBP3 | HALO TAL1 | HALO LYL1 |
|---|---|---|---|---|---|---|---|---|
| ATXN3 | YES | N.D. | | | | | | |
| JOSD1 | YES | NO | | | | | | |

#### USP FAMILY (35 Tested)

| | Present in JURKAT? | ΣΛΟΩ ΓΡΟΩΤΗ? | HALO LMO2 | HALO LDB1 | HALO SSBP2 | HALO SSBP3 | HALO TAL1 | HALO LYL1 |
|---|---|---|---|---|---|---|---|---|
| USP1 | ? | SEVERE | | | YES | | YES | |
| USP2 | NO | NO | | | | | | |
| USP3 | YES | NO | YES | YES | YES | YES | | YES |
| USP4 | YES | NO | YES | YES | YES | YES | | |
| USP5 | YES | SEVERE | | | | | | |
| USP6 | YES | N.D. | | | | | | |
| USP7 | YES | YES | | | | | | |
| USP8 | YES | N.D. | | | | | | |
| USP9X | YES | N.D. | | | | | | |
| USP9Y | ? | N.D. | | | | | | |
| USP10 | YES | N.D. | | | | | | |
| USP11 | YES | N.D. | | | | | | |
| USP12 | ? | NO | | | | | | |
| USP13 | YES | NO | | | | | | |
| USP14 | YES | N.D. | | | | | | |
| USP15 | YES | YES | | | | | | YES |
| USP16 | YES | NO | | | | | | |
| USP17L | ? | N.D. | | | | | | |
| USP18 | ? | NO | | | | | | |
| USP20 | YES | SEVERE | | | | | | |
| USP21 | ? | NO | | | | | | |
| USP22 | YES | NO | | | | | | |
| USP25 | YES | NO | | | | | | |
| USP26 | ? | NO | | | | | | YES |
| USP28 | YES | SEVERE | | | | | | |
| USP29 | ? | NO | | | | | | |
| USP30 | YES | NO | | | | | | |
| USP31 | NO | YES | | | | | | |
| USP32 | YES | SEVERE | | | | | | |
| USP33 | YES | NO | | | | | | |
| USP34 | YES | N.D. | | | | | | |
| USP36 | YES | NO | | | | | | |
| USP37 | YES | NO | | | | | | |
| USP38 | YES | N.D. | | | | | | |
| USP39 | ? | NO | | | | | | |
| USP40 | YES | NO | | | YES | | | YES |
| USP42 | YES | NO | | | | | | |
| USP44 | ? | NO | | | | | | |
| USP46 | NO | NO | | | | | | |
| USP47 | YES | NO | | | | | | |
| USP48 | YES | NO | | YES | YES | YES | | YES |
| USP49 | ? | NO | | | | | | YES |
| USP50 | ? | N.D. | | | | | | |
| USP51 | ? | N.D. | | | | | | |
| USP53 | NO | N.D. | | | | | | |
| USP54 | NO | N.D. | | | | | | |
| USPL1 | ? | NO | | | | | | |
| CYLD | YES | NO | | | | | | |
| PAN2 | ? | YES | | | | | | |
| | | | HALO LMO2 | HALO LDB1 | HALO SSBP2 | HALO SSBP3 | HALO TAL1 | HALO LYL1 |

FIGURE S5

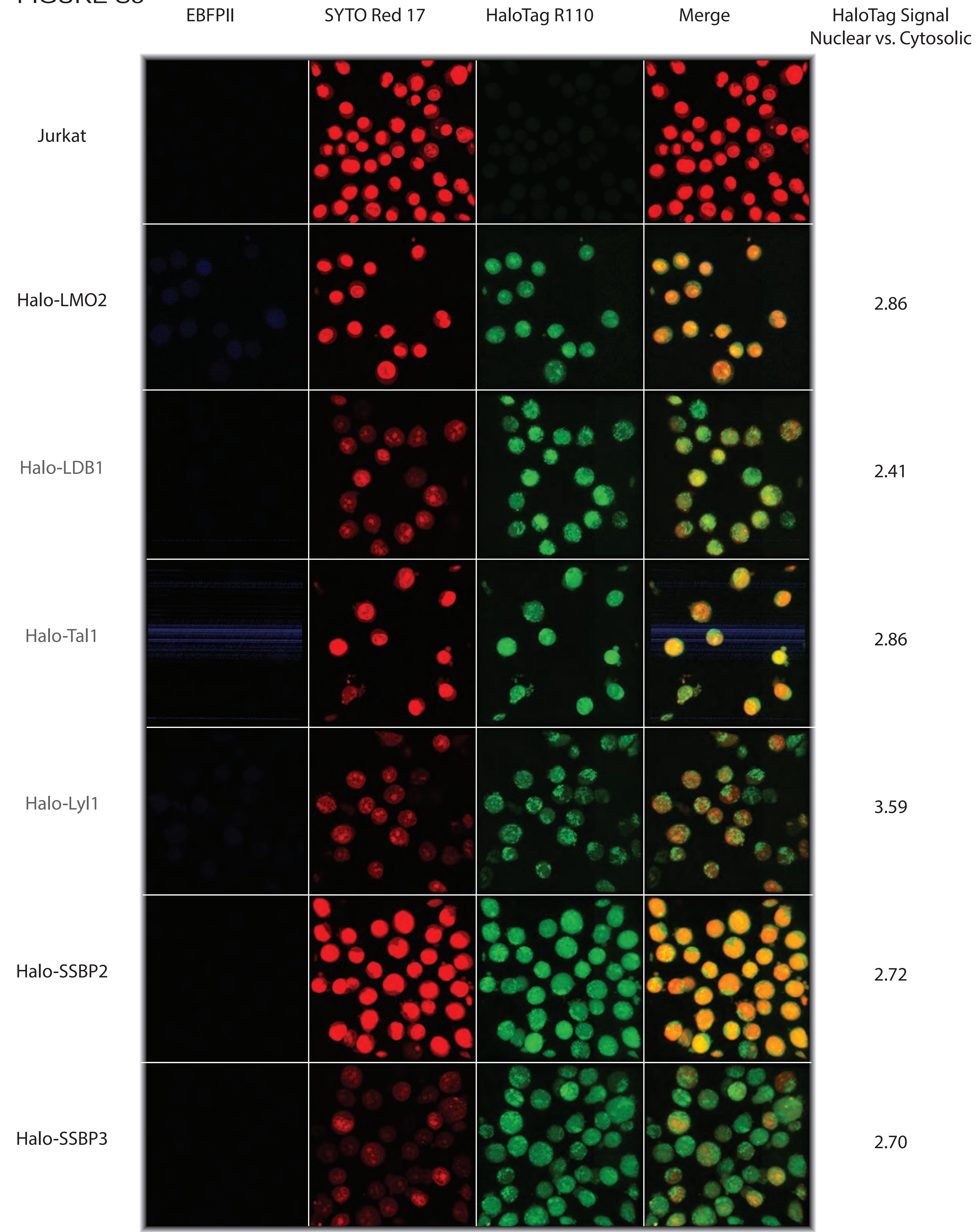
EBFPII
SYTO Red 17
HaloTag R110
Merge
HaloTag Signal
Nuclear vs. Cytosolic
Jurkat
Halo-LMO2
2.86
Halo-LDB1
2.41
Halo-Tal1
2.86
Halo-Lyl1
3.59
Halo-SSBP2
2.72
Halo-SSBP3
2.70
